# Reduced inhibition, bursting, and accelerated oscillations drive early hippocampal hyperactivity in Alzheimer’s disease

**DOI:** 10.1101/2025.06.05.658038

**Authors:** Soraya Meftah, Sungmin Kang, Mingshan Liu, Xingran Wang, Ada Nursel Topçu, Maialen Martin Abad, Áron Kőszeghy, Long Wan, Conor Mullin, Lida Zoupi, Jian Gan

## Abstract

‘Early hippocampal hyperactivity’ is a well-documented yet poorly defined phenomenon in Alzheimer’s disease (AD). While reported in both patients and animal models, its functional manifestations and underlying neurophysiological mechanisms *in vivo* remain unclear.

Here, we address this gap using *in vivo* high-resolution patch-clamp, high-throughput single-unit, and local field recordings in young amyloidopathy mice, at a stage when Aβ remains largely soluble. We uncover previously unidentified cellular mechanisms *in vivo*, characterised by reduced inhibitory synaptic input, hypoactivity of fast-spiking interneurons, and enhanced bursting in pyramidal neurons. At the network level, we reveal accelerated hippocampal oscillations, marked by increased theta and beta power, a departure from the conventional view of oscillation slowing in AD. Mechanistically, this acceleration stems from strengthened synchrony of excitatory currents at higher frequencies and an overall reduction in oscillation-associated inhibitory currents.

Our findings provide the first direct *in vivo* evidence linking early hippocampal hyperactivity to specific synaptic transmission and network dysfunctions, resolving a long-standing ambiguity. Moreover, we propose accelerated oscillations in the hippocampus as a functional biomarker for early AD and a potential therapeutic target for restoring network stability before cognitive decline occurs.

## Introduction

Alzheimer’s disease (AD) is a devastating neurodegenerative condition that profoundly impairs cognitive function and quality of life, imposing significant burdens on individuals, families, and society. Pathologically, AD is characterised by the gradual accumulation of amyloid-beta (Aβ) proteins, transitioning from an early stage of soluble oligomers and fibrils to a late stage of diffuse and compact plaques, alongside intra-and extracellular tau aggregates^1,2^. Recent evidence highlights the toxic role of soluble Aβ in early AD pathophysiology *in vivo*, particularly in the hippocampus^3,4^, with studies suggesting that this toxicity may be reversible^3–5^. Consequently, early detection and intervention for AD are increasingly viewed as both critical and feasible.

Beyond blood and cerebrospinal fluid biomarkers, functional markers aimed at detecting early alterations in neural activity have been investigated, primarily through electroencephalogram (EEG) recordings of neocortical activity, positron emission tomography (PET) and functional magnetic resonance imaging (fMRI) for deeper brain structures^2^. However, technical limitations, particularly in the spatial and temporal resolution, have hindered the exploration of neural activity in deep brain regions such as the hippocampus. The hippocampus plays a pivotal role in memory and spatial cognition—functions that are severely disrupted in AD—and is among the earliest brain regions affected by the disease, even prior to the clinical onset of dementia^6^. Thus, identifying a quantifiable marker of early hippocampal dysfunction would hold immense diagnostic and therapeutic value.

A promising functional marker is early hippocampal hyperactivity^7^. In human studies, this phenomenon has been observed as an enhanced blood oxygen level dependent (BOLD) signal on fMRI in patients with mild cognitive impairment (MCI)^8,9^, presymptomatic familial AD^10^, and carriers of high-risk AD gene, such as APOE4, before clinical AD symptoms^11–14^. Moreover, reducing this early hyperactivity in the hippocampus pharmacologically improved cognitive performance in patients^9^, highlighting its clinical relevance. Similarly, *in vivo* studies of AD mouse models have reported an increased proportion of hyperactive hippocampal neurons via two-photon Ca^2+^ imaging when Aβ is soluble^3^, or upon direct application of Aβ oligmers^4^.

Thus, evidence from human patients and animal models establishes hippocampal hyperactivity as a robust and reproducible feature of early AD. However, the functional manifestations of early hippocampal hyperactivity at the level of network oscillations *in vivo* remain poorly understood, limiting its potential as a clinically quantifiable marker. Furthermore, while early hippocampal hyperactivity has been linked to impaired glutamate reuptake and disruptions in glutamate homeostasis^4^, its cellular mechanisms, and in particular, the contribution of neuronal intrinsic properties and synaptic transmission—both excitatory and inhibitory—remain largely unexplored *in vivo*. Addressing these gaps will be critical for elucidating the mechanisms underlying early hippocampal dysfunction in AD and for developing effective diagnostic and therapeutic strategies.

In this study, we set out to uncover the functional nature of ‘early hippocampal hyperactivity’ by examining early changes in hippocampal oscillations and the underlying synaptic mechanisms *in vivo* in a mouse model of amyloidopathy. Employing *in vivo* patch-clamp and simultaneous local field potential (LFP) recordings in young (2-3 months) APP_swe_/PS1ΔE9 (henceforth APP/PS1) mice^15^, we demonstrated a reduction of basal synaptic inhibition, contributing to overall imbalanced excitation/inhibition inputs. In addition, enhanced phase-locking of synaptic excitation to theta but not delta oscillations underlies accelerated hippocampal oscillations. Furthermore, using high-throughput silicon probe recordings in young (3-5 months) awake animals, we observed similar accelerated hippocampal oscillations characterised by increased beta power. At the cellular level, pyramidal cells exhibited stronger burst firing of action potentials, whereas fast-spiking interneurons showed hypoactivity, as evidenced by increased proportion of low-firing-rate population, consistent with reduced synaptic inhibition revealed by *in vivo* patch-clamp recordings. In summary, our findings provide direct *in vivo* evidence for mechanisms underpinning the long-debated phenomenon of early hippocampal hyperactivity in early amyloidopathy at cellular and functional network levels. We also identify a potential quantifiable functional marker of abnormal neural oscillations in the hippocampus that may aid in early diagnosis.

## Materials and methods

All experiments were carried out in strict accordance with national laws and institutional regulations/guidelines for animal experimentation. Protocols were approved by Home Office project licence (PP8564759).

### *in vivo* patch-clamp and LFP recordings

Male or female 2- to 3-month-old APP/PS1 mice and their littermate wildtype (WT) controls of C57BL/6J background were used. Mice were anesthetised by 100 mg kg^-1^ ketamine and 10 mg kg^-1^ xylazine, and head-fixed in a stereotaxic frame. *In vivo* whole-cell patch-clamp recordings from CA1 pyramidal neurons and simultaneous local field potential (LFP) recordings were performed. Excitatory postsynaptic currents (EPSCs) and inhibitory postsynaptic currents (IPSCs) were recorded in the voltage-clamp configuration with the same cell held at either −70 mV or +10 mV, respectively. Neuronal intrinsic property measurements were performed in separate current-clamp recordings. A glass patch pipette was placed at CA1 pyramidal cell layer of the dorsal hippocampus for LFP recording. LFP signals were down-sampled to 2 kHz, and then filtered between 0.1 Hz and 300 Hz. Power spectrum was obtained by pwelch function in Matlab (window: 2s, overlap: 50%). Absolute power was calculated as the sum of power spectral density within each frequency band. Normalized power for delta (0.1-4Hz), theta (4-12 Hz), and gamma (40–80 Hz) oscillations were calculated from dividing absolute power of each frequency range by total power (0.1∼300Hz), respectively. To quantitatively examine the temporal structure and relationship between synaptic currents and hippocampal oscillations *in vivo*, a derivative-based detection method^16,17^ was used to identify synaptic events. Instantaneous phase of oscillations was calculated by Hilbert transform. Each excitatory postsynaptic current (EPSC) or inhibitory postsynaptic current (IPSC) onset time point was then assigned a Hilbert phase value. Detailed experimental procedures and data analysis for *in vivo* patch-clamp and LFP recordings are provided in the supplementary material.

### High-throughput silicon probe recordings in awake, behaving animals

Male or female 3- to 5-month-old APP/PS1 mice and their littermate WT controls of C57BL/6J background were used. Mice were habituated to head-restraint on an air-cushioned styrofoam ball in a virtual reality (VR) system (JetBall, Phenosys), and trained to run in a VR linear treadmill task for liquid rewards, covering a 150 cm running path per trial, with 10s inter-trial intervals. Silicon probe (A4X32-Poly2–5mm-23s-200-177, NeuroNexus), containing 128 recording channels across four parallel shanks, was slowly inserted into the hippocampus through previously prepared craniotomies. Neural signals were recorded using an Intan 128-channel head-stage (C3316, Intan Technologies) connected to an Intan RHD recording controller (C3004, Intan Technologies). Signals were sampled at 20 kHz and digitised as 16-bit signed integers. Spike sorting was performed using Kilosort 3^18^, followed by manual curation in Phy 2^19^. To classify single-unit spike clusters into putative excitatory and inhibitory neurons, single-units were distinguished based on spike waveform shape and the first moment of the autocorrelogram as described previously^20–23^. Complex bursts were classified as a series of three or more spikes with inter-spike intervals of less than 5 ms. The burst index was defined as the ratio of bursting spikes to all spikes^24^. LFP signals from each site were bandpass filtered (150– 250 Hz), and the channel with the largest mean power of filtered LFP was determined for each shank in each session and designated as the pyramidal cell layer ^25,26^. The power spectral density of neural activity in the pyramidal layer was computed using the Matlab function pwelch (window: 2s; overlap: 50%). Power was calculated as the area under the curve (AUC) of the power spectral density within each frequency band: delta (0.1– 4Hz), theta (4–12 Hz), beta (15– 25 Hz), and gamma (25–75Hz). Detailed experimental procedures and data analysis for high-throughput silicon probe recordings in awake, behaving animals are provided in the supplementary material.

### Statistical analysis

Data are presented as means ± standard error of the mean (SEM). Group comparisons between APP/PS1 and WT genotypes were performed using a rank-based (non-parametric equivalent) generalized linear mixed model (GLMM), where genotype is treated as a fixed effect and animal as a random effect. A rank-based (non-parametric equivalent) general linear model (GLM) was used when replicates were animals. Mann-Whitney U-test was also used to assess differences between groups when specified. Kolmogorov–Smirnov test was used for accumulative distributions. Circular uniformity in phase relationship analysis was examined using Rayleigh test, reported for 10% largest derivative extrema. All statistical analyses were conducted using R or Matlab software. Statistical significance was set at *P <* 0.05. Statistical significance is indicated throughout the paper as follows: * (*P* < 0.05), ** (*P* < 0.01), and *** (*P* < 0.001).

## Results

### Synaptic inhibition is reduced in the hippocampus *in vivo* in young APP/PS1 mice

To investigate synaptic mechanisms underlying early hippocampal hyperactivity in AD, we utilised the well-established APP/PS1 mouse model. Our study focused on a young cohort (2–3 months old), at an age where only soluble Aβ aggregates are present (Supplementary fig.1). APP/PS1 animals and their age-matched WT controls were included, with both male and female mice represented.

We conducted *in vivo* whole-cell patch-clamp recordings from hippocampal CA1 pyramidal cells under ketamine/xylazine general anaesthesia (Fig. 1a and b). Voltage-clamp configuration was employed to precisely measure excitatory and inhibitory synaptic inputs within the same neuron, respectively. Spontaneous EPSCs (sEPSCs) and IPSCs (sIPSCs) were recorded to assess synaptic activity (Fig. 1c).

**Fig. 1.**
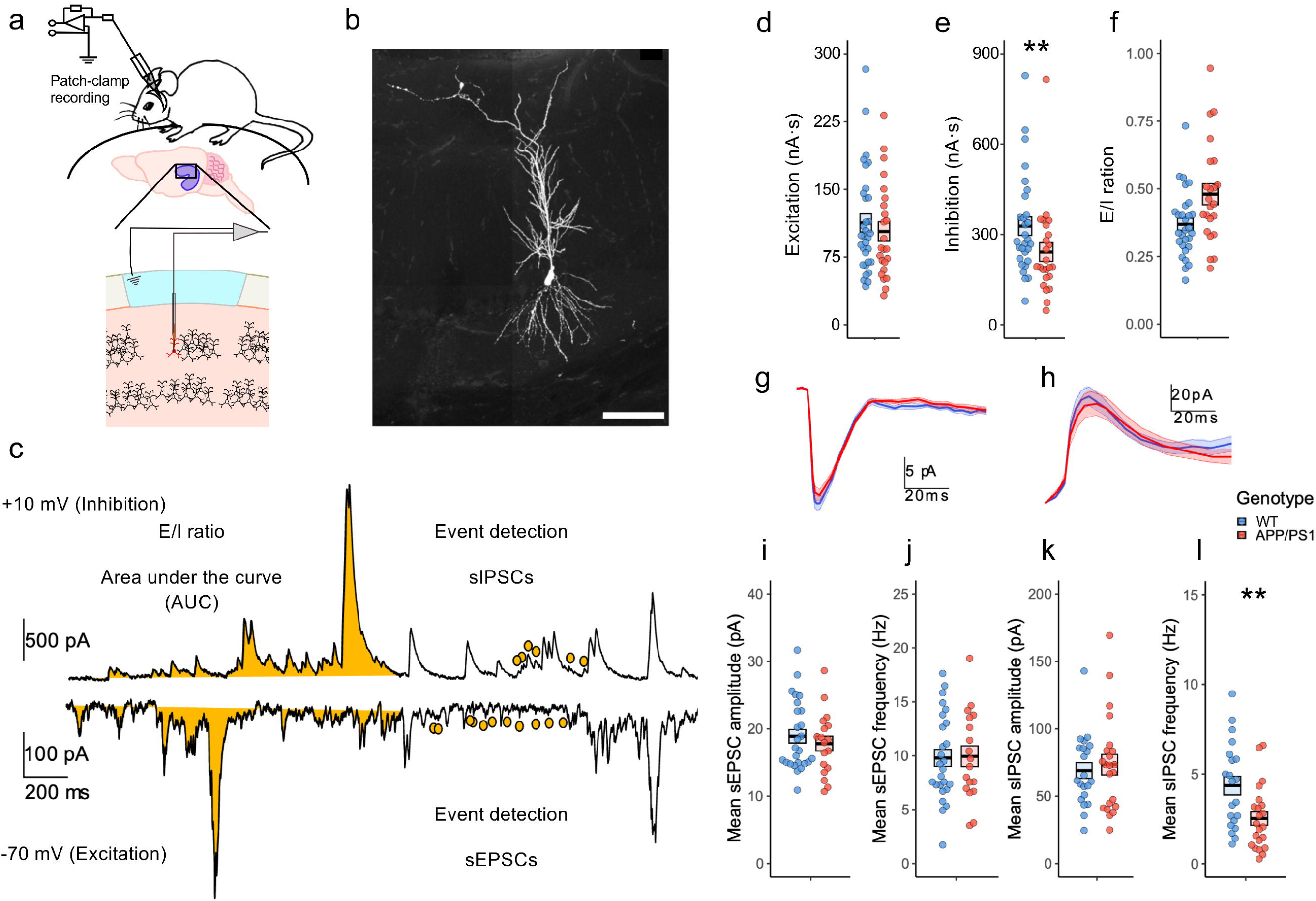
Synaptic excitation and inhibition in young APP/PS1 mice *in vivo*. (**A**) Schematic diagram showing *in vivo* whole-cell patch-clamp recording from the CA1 region of the hippocampus. (**B**) Representative image of a recorded CA1 pyramidal neuron following *in vivo* whole-cell patch-clamp recording. Scale bar: 50 µm. (**C**) Example voltage-clamp traces of spontaneous excitatory (sEPSCs, bottom, −70 mV) and inhibitory (sIPSCs, top, +10 mV) postsynaptic currents. Total charge transfer (area under the curve, AUC, in yellow) was quantified for excitatory and inhibitory current respectively. Filled dots in yellow indicate quantified spontaneous excitatory or inhibitory events using algorithm detection described in methods section. (**D–F**, **I-L**) Box and dot plots show the mean (think line) and SEM (box edges), and each dot represents a single cell. (**D-F**) Overall strength of synaptic excitation, inhibition, and E/I ratio. (**D**) No significant difference in total charge transfer was observed between WT (blue) and APP/PS1 (red) mice. (**E**) APP/PS1 mice exhibited a significant reduction in total inhibitory strength compared to WT mice (*P* = 0.0056). (**F**) E/I ratio was elevated in APP/PS1 mice, though this difference did not reach statistical significance. (**G–H**) Averaged synaptic currents at −70 mV for excitation (sEPSCs) (**G**) and +10 mV for inhibition (sIPSCs) (**H**). Thick line represents the mean and shading represents SEM. (**I–L**) Synaptic transmission properties. (**I**) Mean sEPSC amplitude was comparable between genotypes. (**J**) Mean sEPSC frequency did not differ significantly between groups. (**K**) Mean sIPSC amplitude was largely unchanged. (**L**) Mean sIPSC frequency was significantly reduced in APP/PS1 mice compared to WT (*P* = 0.001), indicating a presynaptic mechanism underlying reduced inhibitory transmission. Asterisks indicate statistical significance (* *P* < 0.05, ** *P* < 0.01).

First, to evaluate the overall strength of synaptic excitation and inhibition, we quantified the total charge transfer (area under the curve). No significant differences in excitatory input strength were observed between genotypes (Fig. 1d; 103.346 ± 10.794 nA·s for APP/PS1, n=24 (14 animals); 112.809 ± 10.041 nA·s for WT, n=32 (18 animals); *P*=0.4258 χ ^2^_(1,56)_ = 0.6343). However, inhibitory input strength was significantly reduced in APP/PS1 mice (Fig. 1e; 241.555 ± 31.078 nA·s for APP/PS1, n=24 (14 animals); 327.758 ± 30.727 nA·s for WT, n=29 (18 animals); *P* =0.0056, χ ^2^_(1,53)_ = 7.6716), but the excitation/inhibition (E/I) ratio remained largely unchanged. (Fig. 1f; 0.4799 ± 0.0387 for APP/PS1, n=23 (14 animals); 0.3692 ± 0.0227 for WT, n=29 (18 animals); *P* = 0.3185, χ ^2^_(1,52)_ = 0.9949).

Next, to investigate potential pre- or postsynaptic mechanisms underlying these changes, we examined the frequency and amplitude of baseline sEPSCs and sIPSCs, respectively. No significant differences were observed in sEPSC amplitude (Fig. 1i; 17.8008 ± 1.1014 pA for APP/PS1, n=18 (12 animals); 18.8982 ± 1.0172 pA for WT, n=26 (17 animals); P =0.8508, χ ^2^_(1,44)_ = 0.0354) or sEPSC frequency (Fig. 1j; 9.9491 ± 0.9667 Hz for APP/PS1, n=18 (12 animals); 9.7949 ± 0.7886 Hz for WT, n=26 (17 animals); P =0.8245, χ ^2^_(1,44)_ = 0.0492). However, a significant reduction in sIPSC frequency (Fig. 1l; 2.5167 ± 0.3794 Hz for APP/PS1, n=22 (13 animals); 4.3531 ± 0.5157 Hz for WT, n=21 (15 animals); *P* = 0.0001, χ ^2^_(1,43)_ = 10.759) was detected, whereas sIPSC amplitude remained overtly unchanged (Fig. 1k; 73.3805 ± 7.6386 pA for APP/PS1, n=22 (13 animals); 69.0283 ± 5.7638 pA for WT, n=21 (15 animals); P =0.0518, χ ^2^_(1,43)_ = 3.7813). This suggested a presynaptic mechanism could be responsible for the overall reduction of inhibitory transmission strength.

To assess potential changes in the intrinsic properties of pyramidal neurons in the CA1 region, we conducted current-clamp recordings in a separate set of CA1 pyramidal cells *in vivo* (Fig. 2a and b). No significant differences were observed in key passive membrane properties, including resting membrane potential (Fig. 2d, −62.0769 ± 1.8483 mV for APP/PS1, n=13 (11 animals);-60.3125 ± 1.7835 mV for WT, n=16 (15 animals); *P* = 0.4686, χ ^2^_(1,29)_ = 0.5253) and input resistance (Fig. 2e, 122.2308 ± 9.7125 MΩ for APP/PS1, n=13 (11 animals); 108.5000 ± 5.4345 MΩ for WT, n=16 (15 animals); *P* = 0.1761, χ ^2^_(1,29)_ = 1.8299). Similarly, no change in input-frequency firing patterns was detected (Fig. 2c). Furthermore, actional potential threshold (Fig. 2f, −44.8846 ± 1.4677 mV for APP/PS1, n=13 (11 animals); −45.9563 ± 1.0392 mV for WT, n=16 (15 animals); *P* = 0.9816, χ ^2^_(1,29)_ = 0.0005), peak amplitude (Fig. 2g, 94.6000 ± 3.3031 mV for APP/PS1, n=13 (11 animals); 98.3063 ± 2.8204 mV for WT, n=16 (15 animals); *P* = 0.3824, χ ^2^_(1,29)_ = 0.7629), and maximal rise speed (Fig. 2h, 694.2462 ± 43.0059 mV·ms^-1^ for APP/PS1, n=13 (11 animals); 739.2437 ± 57.9287 mV·ms^-1^ for WT, n=16 (15 animals); *P* = 0.4329, χ ^2^_(1,29)_ = 0.6151) remained indistinguishable between genotypes. Notably, AP half-width slightly reduced in APP/PS1 animals (Fig. 2i, 0.8054± 0.0200 ms for APP/PS1, n=13 (11 animals); 0.8881 ± 0.0314 ms for WT, n=16 (15 animals; *P* = 0.0459, χ ^2^_(1,29)_ = 3.9838). Overall, these data suggested there was no overt change in intrinsic properties in CA1 pyramidal cells in young APP/PS1 mice *in vivo*.

**Fig. 2.**
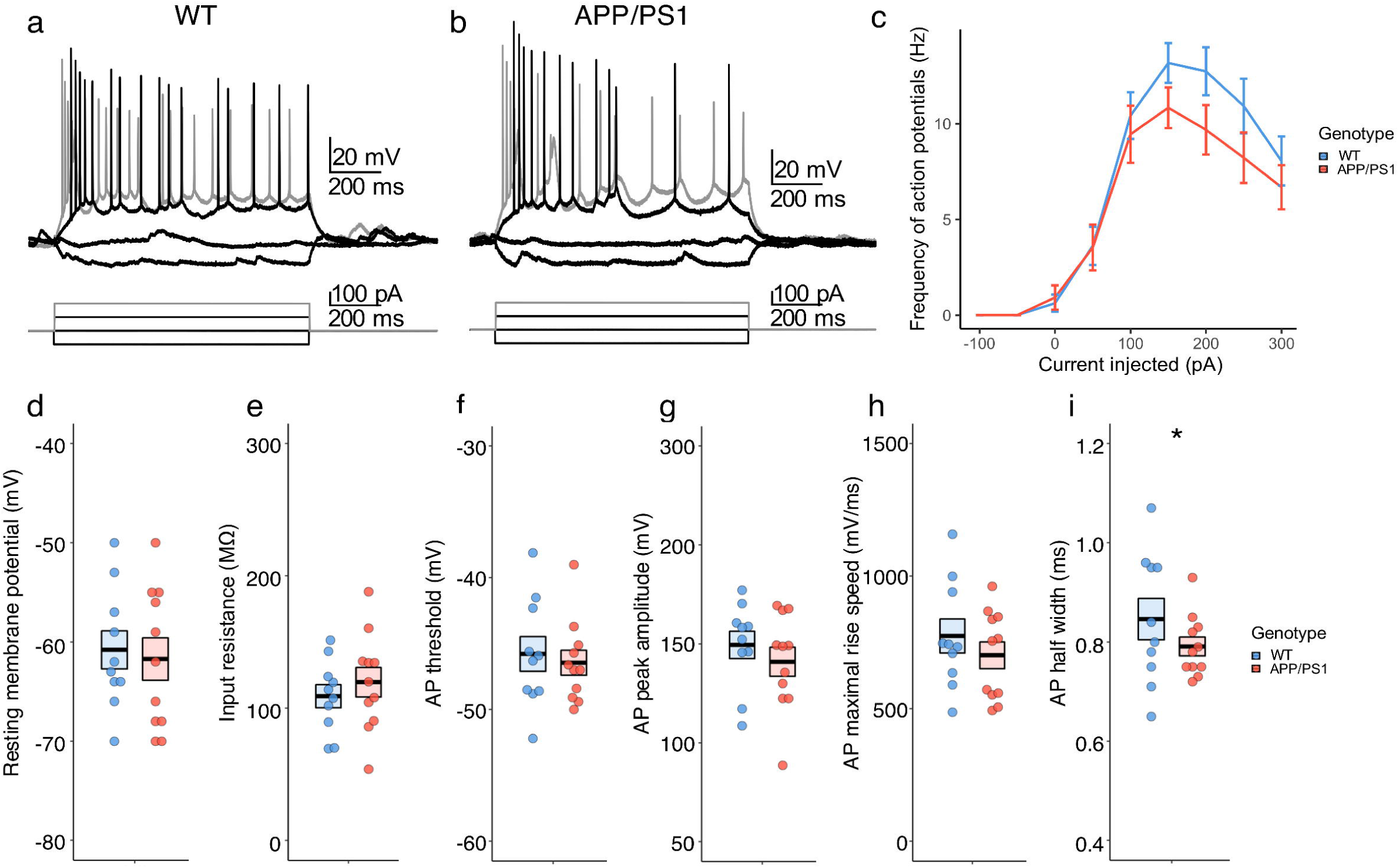
Intrinsic properties of CA1 pyramidal neurons in young APP/PS1 mice *in vivo*. (**A**, **B**) Representative *in vivo* current-clamp recording traces from CA1 pyramidal neurons in WT (**A**) and APP/PS1 (**B**) mice in response to injected stepwise current injections. (**C**) Input-frequency relationship showing action potential frequency as a function of injected current. No significant difference was detected between genotypes. (**D–I**) Quantification of intrinsic membrane and action potential properties. (**D**) Resting membrane potential was comparable between groups. (**E**) Input resistance did not differ significantly between genotypes. (**F**) Action potential threshold remained unchanged. (**G**) Action potential peak amplitude was similar between WT and APP/PS1 mice. (**H**) Maximum action potential rise speed was not significantly different. (**I**) Action potential half-width was significantly increased in APP/PS1 mice compared to WT (*P* = 0.0459), suggesting altered spike kinetics. Asterisks indicate statistical significance (* *P* < 0.05, ** *P* < 0.01).

### Neural oscillations are accelerated in the hippocampus in young APP/PS1 mice *in vivo*

Next, we moved up our focus from single-cell physiology to neural network-level analysis. Neural oscillations are quantifiable measurements of brain function and directly underlie normal cognitive computation. We took advantage of our *in vivo* preparation which preserved not only an intact neural circuitry, but also key oscillations that drive memory encoding, consolidation, and retrieval in the hippocampus. We sought to determine whether hippocampal oscillations are altered in the early stages of AD where Aβ burden is still largely soluble, and if so, to identify the underlying synaptic mechanisms.

To address this, we performed simultaneous LFP and voltage-clamp recordings in young APP/PS1 mice (Fig. 3a and b). We first analysed the power spectrum of recorded LFPs. While no significant differences were observed in total LFP power between genotypes (0.1-300 Hz; Fig. 3d, 0.0642 ± 0.0045 mV^2^ for APP/PS1, n=15 animals; 0.0767 ± 0.0045 mV^2^ for WT, n=19 animals; *P* = 0.0928, F^2^_(1,34)_ = 3.0014), an analysis of the relative power (Fig. 3c) revealed a significant decrease in the delta band (0.1-4Hz; Fig. 3e; 0.8031 ± 0.0152 for APP/PS1, n=15 animals; 0.8484 ± 0.0076 for WT, n=19 animals; *P* = 0.0108, F^2^_(1,34)_ = 7.3297), and a significant increase in the theta band (4-12Hz; Fig. 3f, 0.1208 ± 0.0102 for APP/PS1, n=15 animals; 0.0902 ± 0.0046 for WT, n=19 animals; *P* = 0.0067, F^2^_(1,34)_ = 8.3939) in APP/PS1 animals. The relative power of gamma oscillations remained unchanged (30-100 Hz, Fig. 3g, 0.0205 ± 0.0016 for APP/PS1, n=15 animals; 0.0183 ± 0.0012 for WT, n=19 animals; *P* = 0.2813, F^2^_(1,34)_ = 1.2010). These results characterised by an increased proportion of theta frequency component over delta in LFP, indicated an overall acceleration of hippocampal oscillations in young APP/PS1 mice.

**Fig. 3.**
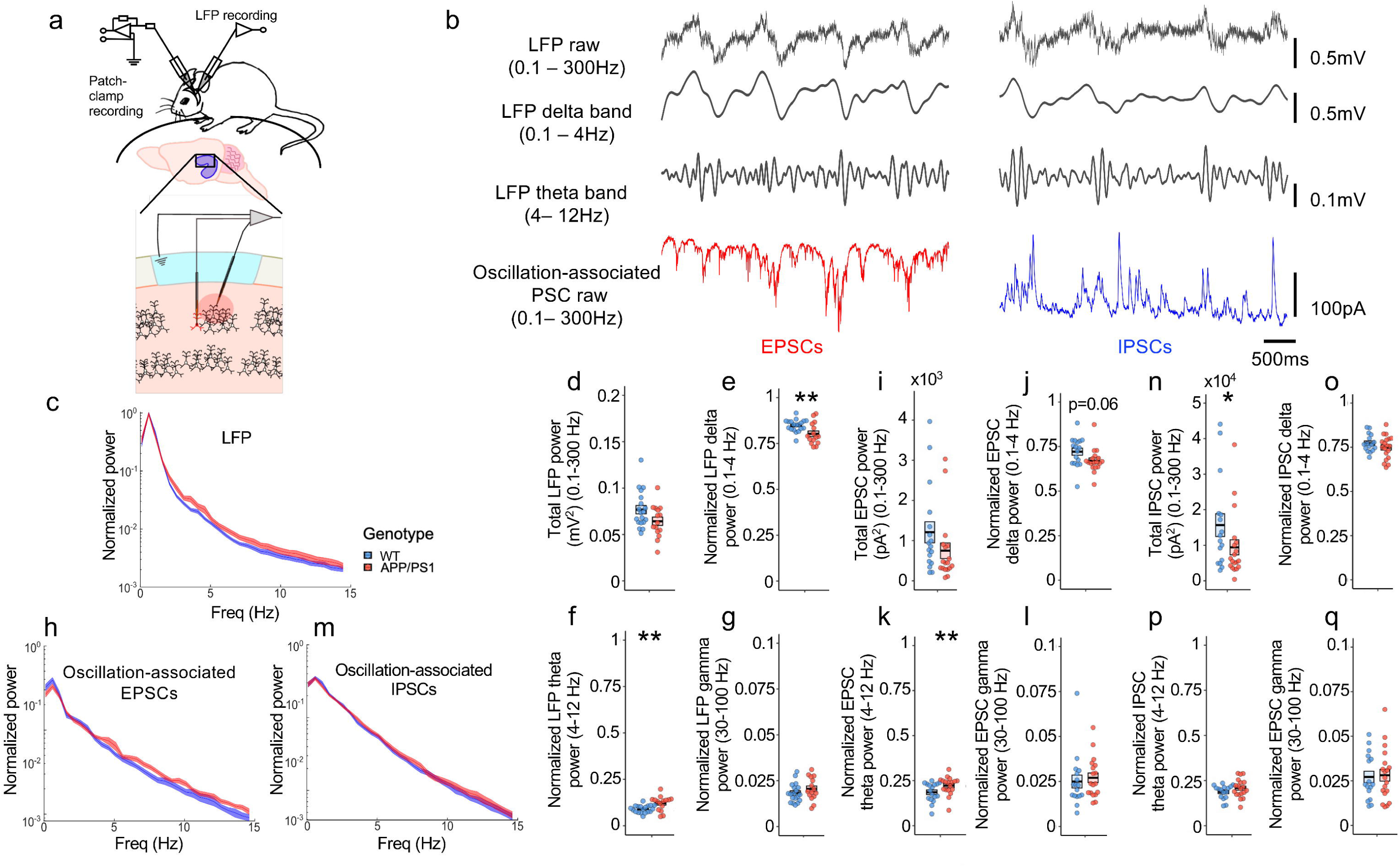
Altered hippocampal network oscillations and associated synaptic currents in young APP/PS1 mice. (**A**) Schematic diagram showing simultaneous *in vivo* voltage-clamp and LFP recordings from the CA1 region of the hippocampus. (**B**) Representative LFP traces (0.1–300 Hz), delta band (0.1–4 Hz), theta band (4–12 Hz), and oscillation-associated postsynaptic currents (PSCs) recorded in WT and APP/PS1 mice. Excitatory (EPSCs, red) and inhibitory (IPSCs, blue) postsynaptic currents are shown. (**C**) Normalized LFP power spectral density in WT (blue) and APP/PS1 (red) mice. (**D–G**) Quantification of total LFP power (**D**) and normalized power in delta (**E**), theta (**F**), and gamma (**G**) frequency bands, respectively. APP/PS1 mice exhibited significantly reduced delta power (*P* = 0.0108) and increased theta power (*P* = 0.0067), indicating an acceleration of hippocampal oscillations. Gamma power remained unchanged (*P* = 0.2813). (**H–M**) Normalized power spectral density of oscillation-associated EPSCs (**H**) and IPSCs (**M**) in WT and APP/PS1 mice, respectively. (**I–L**) Quantification of total EPSC power (**I**) and normalized power in delta (**J**), theta (**K**), and gamma (**L**) frequency bands, revealing a significant increase in theta power in APP/PS1 mice (*P* = 0.006). (**N–Q**) Quantification of total IPSC power (**N**) and normalized power in delta (**O**), theta (**P**), and gamma (**Q**) bands, revealing a significant reduction in total power in APP/PS1 mice (*P* = 0.038). WT mice are shown in blue, APP/PS1 mice in red. Asterisks indicate statistical significance (* *P* < 0.05, ** *P* < 0.01).

### Accelerated hippocampal oscillations associate with phase-locked excitatory transmission at higher oscillatory frequency

LFPs predominantly reflect the summation of synaptic transmission arising from coordinated firings of large neuronal populations. To investigate the synaptic mechanisms underlying the observed acceleration of hippocampal oscillations, we examined oscillation-associated excitatory and inhibitory synaptic transmission that may contribute to the reduction of delta and the increase of theta power in LFPs in APP/PS1 mice.

We first analysed the power spectrum of oscillation-associated synaptic currents. In EPSCs, the total power remained largely unchanged (Fig. 3i; 750.5450 ± 194.9726 pA^2^ for APP/PS1, n=18 (12 animals); 1208.2690 ± 262.4596 pA^2^ for WT, n=17 (14 animals); *P* = 0.1250, χ ^2^_(1,35)_ = 2.3531). When relative power was analysed (Fig. 3h), a decreased trend in the delta frequency (0.1-4Hz) component (Fig. 3j; 0.6706 ± 0.0157 for APP/PS1, n=18 (12 animals); 0.7214 ± 0.0199 for WT, n=17 (14 animals); *P* = 0.0641, χ ^2^_(1,35)_ = 3.4277) and a significant increase in the theta frequency (4-12 Hz) component (Fig. 3k; 0.2225 ± 0.0118 for APP/PS1, n=18 (12 animals); 0.1855 ± 0.0127 for WT, n=17 (14 animals); *P* = 0.0061, χ ^2^_(1,35)_ = 7.5302) were observed. There was no change for gamma frequency (Fig. 3l; 0.0271 ± 0.0027 for APP/PS1, n=18 (12 animals); 0.0250 ± 0.0036 for WT, n=17 (14 animals); *P* = 0.1999, χ ^2^_(1,35)_ = 1.6429). These findings suggested though overall magnitude of oscillation-associated excitatory drive did not change, its inner dynamics became faster in young APP/PS1 mice, which was consistent with the accelerated hippocampal oscillations observed previously.

In contrast, for IPSCs, a significant reduction in total power was observed in APP/PS1 animals (Fig. 3n; 9423.5340 ± 2123.5060 pA^2^ for APP/PS1, n=19 (12 animals); 15676.2360 ± 3210.7490 pA^2^ for WT, n=16 (14 animals); *P* = 0.0383, χ ^2^_(1,35)_ = 4.2911), but no change was observed in relative power (Fig. 3m) at delta (0.1- 4Hz; Fig. 3o; 0.7492 ± 0.0152 for APP/PS1, n=19 (12 animals); 0.7735 ± 0.0124 for WT, n=16 (14 animals); *P* = 0.2489, χ ^2^_(1,35)_ = 1.3297), theta (4-12 Hz; Fig3p; 0.2080 ± 0.0117 for APP/PS1, n=19 (12 animals); 0.1871 ± 0.0097 for WT, n=16 (14 animals); *P* = 0.4928, χ ^2^_(1,35)_ = 0.4728), and gamma (30-100Hz; Fig3q; 0.0056 ± 0.0007 for APP/PS1, n=19 (12 animals); 0.0054 ± 0.0006 for WT, n=16 (14 animals); *P* = 0.9301, χ ^2^_(1,35)_ = 0.0077) frequency ranges.

Based on these findings, we further examined the precise temporal relationship between excitatory or inhibitory synaptic events with hippocampal oscillations in young APP/PS1 mice by performing a phase-locking analysis. We first detected individual EPSCs and IPSCs occurring in conjunction with hippocampal oscillations using a derivative-based event detection method^16,17^, and then analysed their timing relative to oscillation cycles of delta (0.1-4Hz) and theta (4-12 Hz) bands (Fig. 4a, see details in method). Specifically, we plotted the timing of these events against the phase space of the oscillation cycle. Our analysis revealed that EPSCs were phase-locked to both delta (Fig. 4b; 96.7° ± 22.1°, n=18, *P* = 0.0430) and theta (Fig. 4c; 99.7° ± 18.5°, n=29; *P* = 0.0032) oscillations in APP/PS1 animals, but less so to delta compared to theta in WT animals (For delta: 81.0° ± 22.5°, n=17, *P* = 0.2115; For theta: 86.8° ± 21.0°, n=17, *P* = 0.0033). IPSCs however, were phase-locked to both delta and theta oscillations across genotypes (Fig. 4e for delta; 126.5° ± 19.4° for APP/PS1, n=19, *P* = 0.0002; 95.0 ± 9.5 for WT, n=16, *P* = 0.0011; Fig. 4f for theta; 95.6° ± 9.7° for APP/PS1, n=19, *P* < 0.0001; 101.5 ± 12.7 for WT, n=16, *P* = 0.0001). By quantifying the strength of phase-locking through vector length calculations, we observed significantly stronger phase-locking of EPSCs to theta (Fig. 4d Right; 0.9482 ± 0.0179 for APP/PS1, n=18 (12 animals); 0.8266 ± 0.0372 for WT, n=17 (14 animals); *P* =0.0006, χ ^2^_(1,35)_ = 11.5650), but not delta (Fig. 4d Left; 0.7915 ± 0.0434 for APP/PS1, n=18 (12 animals); 0.7802 ± 0.0547 for WT, n=17 (14 animals); *P* = 0.7946, χ ^2^_(1,35)_ = 0.0678) oscillations in young APP/PS1 mice. In contrast, IPSCs did not show difference in their phase-locking, either to delta (Fig. 4g Left; 0.6884 ± 0.0536 for APP/PS1, n=19 (12 animals); 0.6676 ± 0.0669 for WT, n=16 (14 animals); *P* = 0.922, χ ^2^_(1,35)_ =0. 0096), or theta (Fig. 4g Right; 0.9138 ± 0.0255 for APP/PS1, n=19 (12 animals); 0.8423 ± 0.0465 for WT, n=16 (14 animals); *P* = 0.2414, χ ^2^_(1,35)_ = 1.3727) oscillations. These findings suggested an enhanced synchrony between excitatory synaptic events and hippocampal theta oscillations in young APP/PS1 mice, supporting the findings from prior power spectrum analysis (Fig. 3). Our results indicated that the observed acceleration of hippocampal oscillations in young APP/PS1 mice could be attributed to a selectively elevated synchrony of excitatory synaptic drive arriving at a higher frequency.

**Fig. 4.**
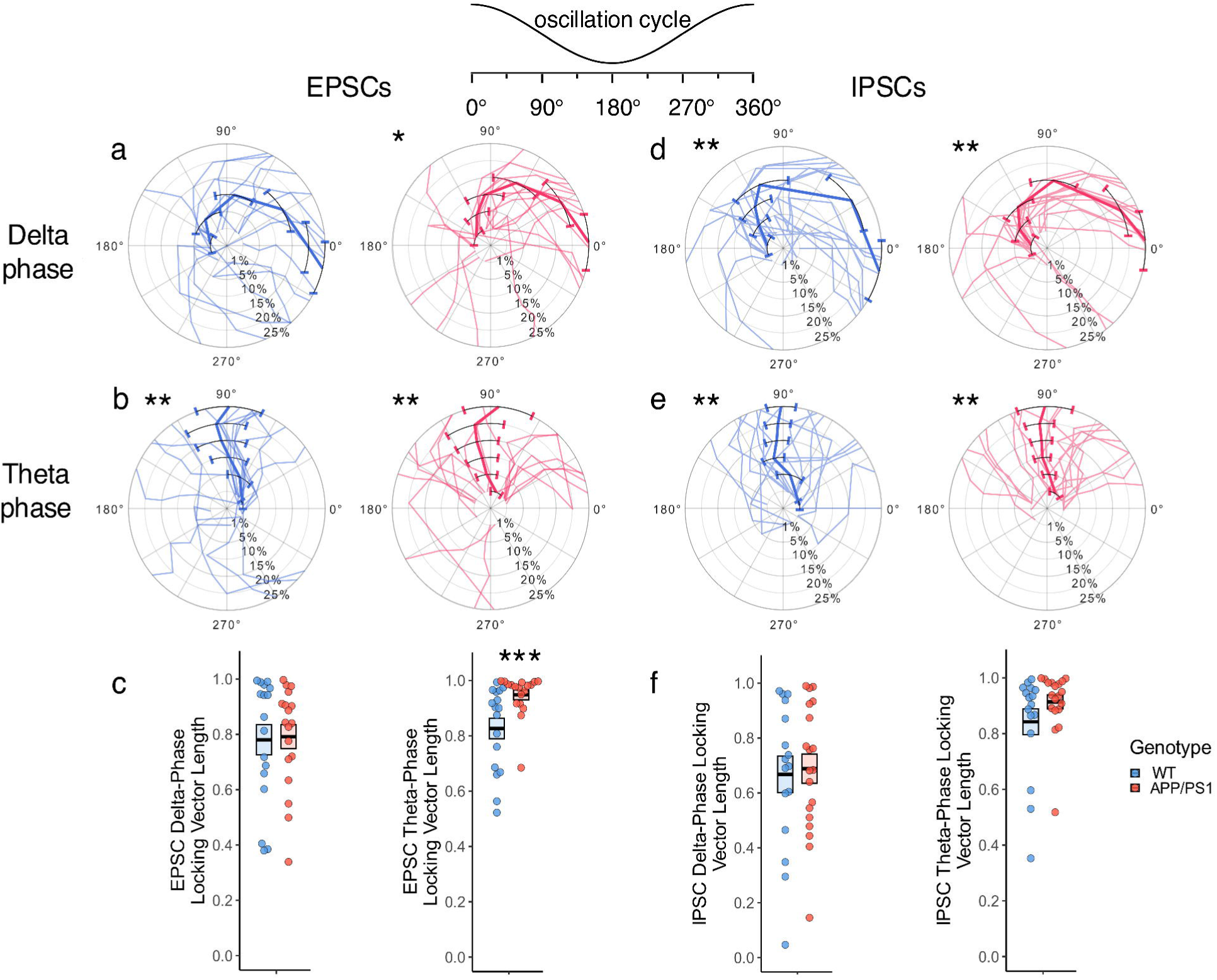
Phase-locking of excitatory and inhibitory synaptic currents to hippocampal oscillations in young APP/PS1 mice. (**A**, **B**) Polar plots of mean EPSC onsets relative to delta (A, 0.1–4 Hz) and theta (B, 4–12 Hz) oscillations in WT (blue) and APP/PS1 (red) mice. Thin lines represent individual cell data, thick coloured lines and symbols indicate mean values with angular deviation. Non-uniformity was tested using Rayleigh test for 10% largest derivative peaks. (**C**) Quantification of vector strength for EPSC phase-locking to delta and theta oscillations. APP/PS1 mice showed significantly increased EPSC phase-locking to theta oscillations (*P* = 0.0006). (**D**, **E**) Same as A and B but for IPSC onsets. (**F**) Quantification of vector strength for IPSC phase-locking to delta and theta oscillations. No significant differences were detected between groups. Asterisks indicate statistical significance (* *P* < 0.05, ** *P* < 0.01, *** *P* < 0.001).

### Hypoactivity of hippocampal fast-spiking interneurons in awake young APP/PS1 mice

To avoid potential confounding effects of general anaesthesia on cellular physiology and network-level properties, we sought to validate our findings in drug-free, behaving animals. Head-restrained young APP/PS1 mice (3–5 months old) were trained to run on a linear treadmill within a virtual reality environment (Fig. 5a). Using high-density silicon probe recordings, we simultaneously captured single-neuron activity and LFPs in the hippocampus during task performance (Fig. 5b). In total, we recorded 757 neurons from 34 recording sessions in 7 APP/PS1 mice and 975 neurons from 40 recording sessions in 9 WT controls.

**Fig. 5.**
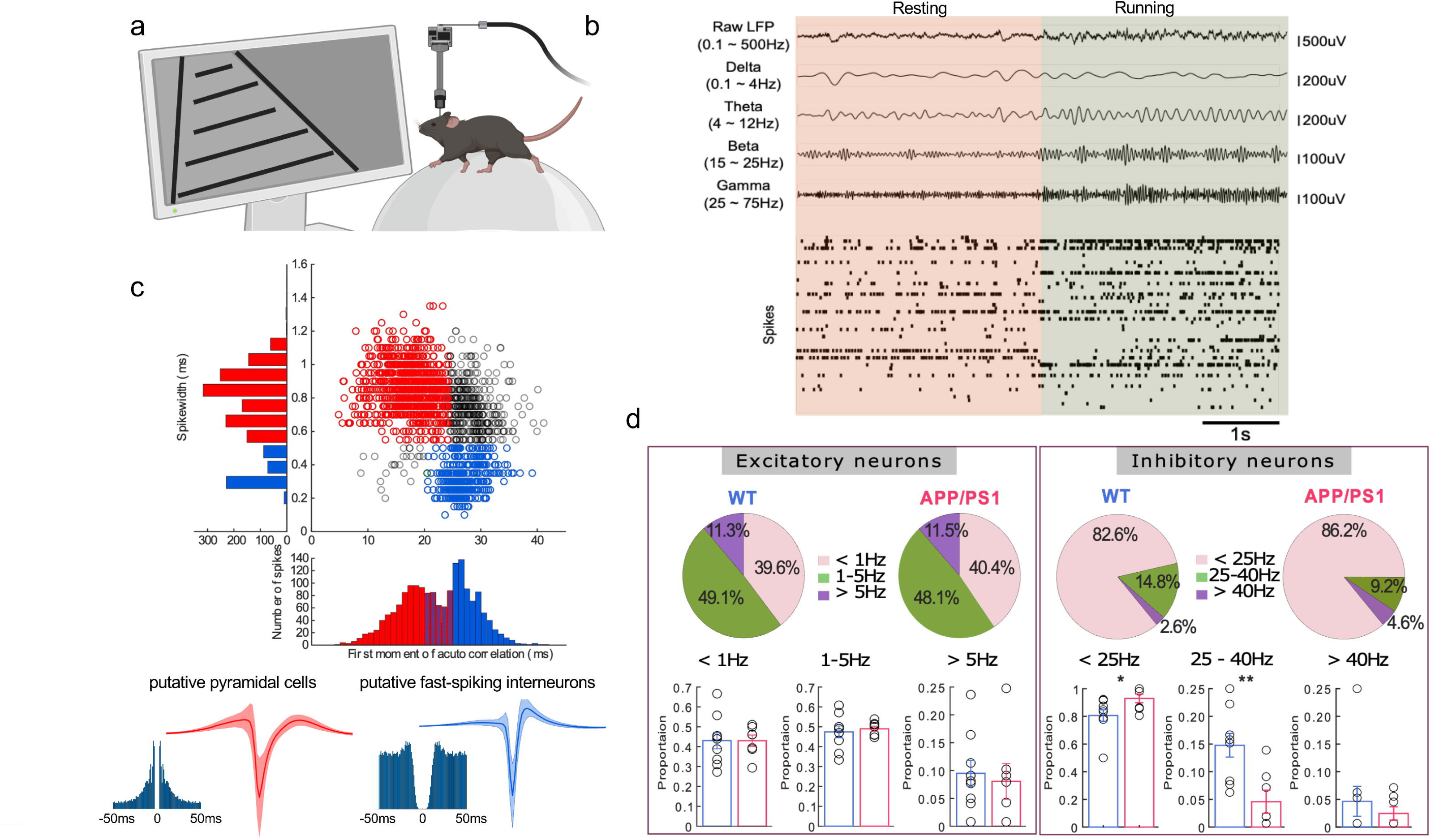
Firing properties of hippocampal neurons in awake behaving young APP/PS1 mice. (**A**) Schematic diagram showing high-throughput silicon probe recording from awake, behaving mice in a virtual reality (VR) system. (**B**) Representative LFPs and spike times during resting (*left*, pink background) and running (*right*, green background) conditions. In raster plots, each row of the raster represents an individual neuron, each vertical tick represents the time of an action potential. (**C**) Classification of putative pyramidal cells (red) and fast-spiking interneurons (blue) based on spike width and first moment of autocorrelogram properties. Below, representative autocorrelograms and average waveforms with standard deviation (shaded) are shown for each cell type. (**D**) Distribution of firing rates in CA1 neurons. (*Left*) Pie charts showing the proportion of pyramidal cell populations of different firing rates (<1 Hz, 1–5 Hz, >5 Hz). Proportions were quantified per animal, shown in bar charts. No significant genotype difference was found. (*Right*) Pie charts showing the proportion of fast-spiking interneuron populations of different firing rates (<25 Hz, 25–40 Hz, >40 Hz). Proportions were quantified per animal, shown in bar charts. Note an increased proportion of low-firing (<25 Hz, *P* = 0.023) and a decreased proportion of medium-firing fast-spiking interneurons (25–40 Hz, *P* = 0.0004) in APP/PS1 mice. High-firing proportion remained unchanged. Asterisks indicate statistical significance (* *P* < 0.05, ** *P* < 0.01).

We initiated our analysis at the single-cell level to investigate whether hippocampal pyramidal cells and interneurons are differentially affected by the accumulation of soluble Aβ. To this end, we identified putative pyramidal cells and fast-spiking interneurons from well-clustered single units using established criteria^20–23^ (see details in methods). This classification resulted in the identification of 948 putative pyramidal cells, 264 putative fast-spiking interneurons, and 520 unclassified units (Fig. 5c).

We first analysed the general firing profiles of hippocampal neurons when the animal engaged with the linear treadmill task. The proportion of putative pyramidal cells was comparable across low (<1 Hz, 0.4297 ± 0.0289 for APP/PS1, n = 7 animals; 0.4306 ± 0.0413 for WT, n = 9 animals; *P* = 0.6135, f^2^_(1,16)_ = 0.2668), medium (1–5 Hz, 0.4900 ± 0.0131 for APP/PS1, n = 7 animals; 0.4745 ± 0.0295 for WT, n = 9 animals; *P* = 0.6490, f^2^_(1,16)_ = 0.2163), and high (>5 Hz, 0.0804 ± 0.0320 for APP/PS1, n = 7 animals; 0.0949 ± 0.0235 for WT, n = 9 animals; *P* = 0.6858, f^2^_(1,16)_ = 0.1706) firing rate ranges between WT and APP/PS1 mice (Fig. 5d Left). In contrast, we observed notable changes in firing rate distributions in putative fast-spiking interneurons. Specifically, the proportion of low-firing-rate population (<25 Hz) was significantly increased in APP/PS1 animals compared to WT controls (0.9296 ± 0.0318 for APP/PS1, n = 7 animals; 0.8059 ± 0.0422 for WT, n = 9 animals; *P* = 0.023, f^2^_(1,16)_ = 6.4749), whereas the proportion of medium-firing-rate population (25–40 Hz) was decreased (0.0458 ± 0.0209 for APP/PS1, n = 7 animals; 0.1477 ± 0.0214 for WT, n = 9 animals; *P* = 0.0004, f^2^_(1,16)_ = 11.533). The proportion of high-firing-rate population (>40 Hz) did not differ significantly between genotypes (0.0246 ± 0.0120 for APP/PS1, n = 7 animals; 0.0464 ± 0.0270 for WT, n = 9 animals; *P* = 0.8679, f^2^_(1,16)_ = 0.0287) (Fig. 5d Right). These data indicated an overall ‘hypoactivity’ of fast-spiking interneurons in drug-free, task-engaging, young APP/PS1 mice. Given that fast-spiking interneurons account for about 80% of total interneuron populations in the hippocampus, these results could explain the previous finding of an overall reduction in basal inhibitory transmission from *in vivo* patch-clamp recording (Fig. 1e and i).

### Enhanced bursting of CA1 pyramidal cells in awake young APP/PS1 mice

Hyperactivity of CA1 pyramidal cells in young APP/PS1 mice has been observed by *in vivo* 2-photon Ca^2+^ imaging^3^. However, the inadequate temporal resolution offered by Ca^2+^ indicators may have hampered unequivocal discrimination of single spikes versus bursts *in vivo*, leaving the cellular correlates of ‘hyperactivity’ largely unresolved. In addition, this finding was reported under general anaesthesia. To overcome these limitations, we took advantage of high temporal resolution using high-throughput silicon probe electrophysiology in awake mice.

We analysed action potential firing patterns in putative pyramidal cells from our experiments (Fig. 6a and b). Although the percentage of bursting cells did not significantly differ between groups (WT: 0.7766 ± 0.0290, APP/PS1: 0.7842 ± 0.0437; *P* = 0.7108, W = 27.5), the distribution of inter-spike intervals (ISIs) within bursts revealed that APP/PS1 mice had significantly shorter ISIs during bursting and higher instantaneous bursting frequency compared to WT controls (Fig. 6c; Kolmogorov–Smirnov test, *P* = 0.0026). In addition, there was a trend of increased intra-burst spike number in APP/PS1 animals (Fig. 6d). During resting periods between task runs, burst event rate was comparable between groups (Fig. 6b left; APP/PS1: 0.0515 ± 0.0068, 295 cells from 7 animals; WT: 0.0556 ± 0.0056, 493 cells from 9 animals; χ ^2^_(1,788)_ =0.0168, *P* = 0.8969). Similarly, the burst index showed no significant difference (Fig. 6e left; APP/PS1: 0.0705 ± 0.006, 295 cells from 7 animals; WT: 0.0644 ± 0.0040, 493 cells from 9 animals; χ ^2^_(1,788)_ =0.0491, *P* = 0.8246). However, during active running periods, burst event rate (Fig. 6b right; APP/PS1: 0.0809 ± 0.0126 Hz, 197 cells from 7 animals; WT: 0.0699 ± 0.0076 Hz, 405 cells from 9 animals) and burst index (Fig. 6e right; APP/PS1: 0.0817 ± 0.0078, 197 cells from 7 animals; WT: 0.0725 ± 0.0048, 405 cells from 9 animals) appeared higher in APP/PS1 mice. Mann-Whitney U test showed a significant difference between groups for burst event rate (*P* = 0.0297) and burst index (*P* = 0.0453), but these effects were not significant when using a generalized linear mixed model (GLMM) that accounted for inter-animal variability (burst event rate: χ ^2^_(1,602)_ =1.7203, *P* = 0.1897; burst index: χ ^2^_(1,602)_ =1.659, *P* = 0.1977). This suggested that while there was a trend towards a more frequent bursting activity in APP/PS1 mice during running, it is subtle and variability across animals may contribute to the observed differences. Overall, our findings suggested that young APP/PS1 mice exhibited higher bursting frequency and likely more intra-burst spikes when bursts were generated in hippocampal pyramidal cells than their WT counterparts.

**Fig. 6.**
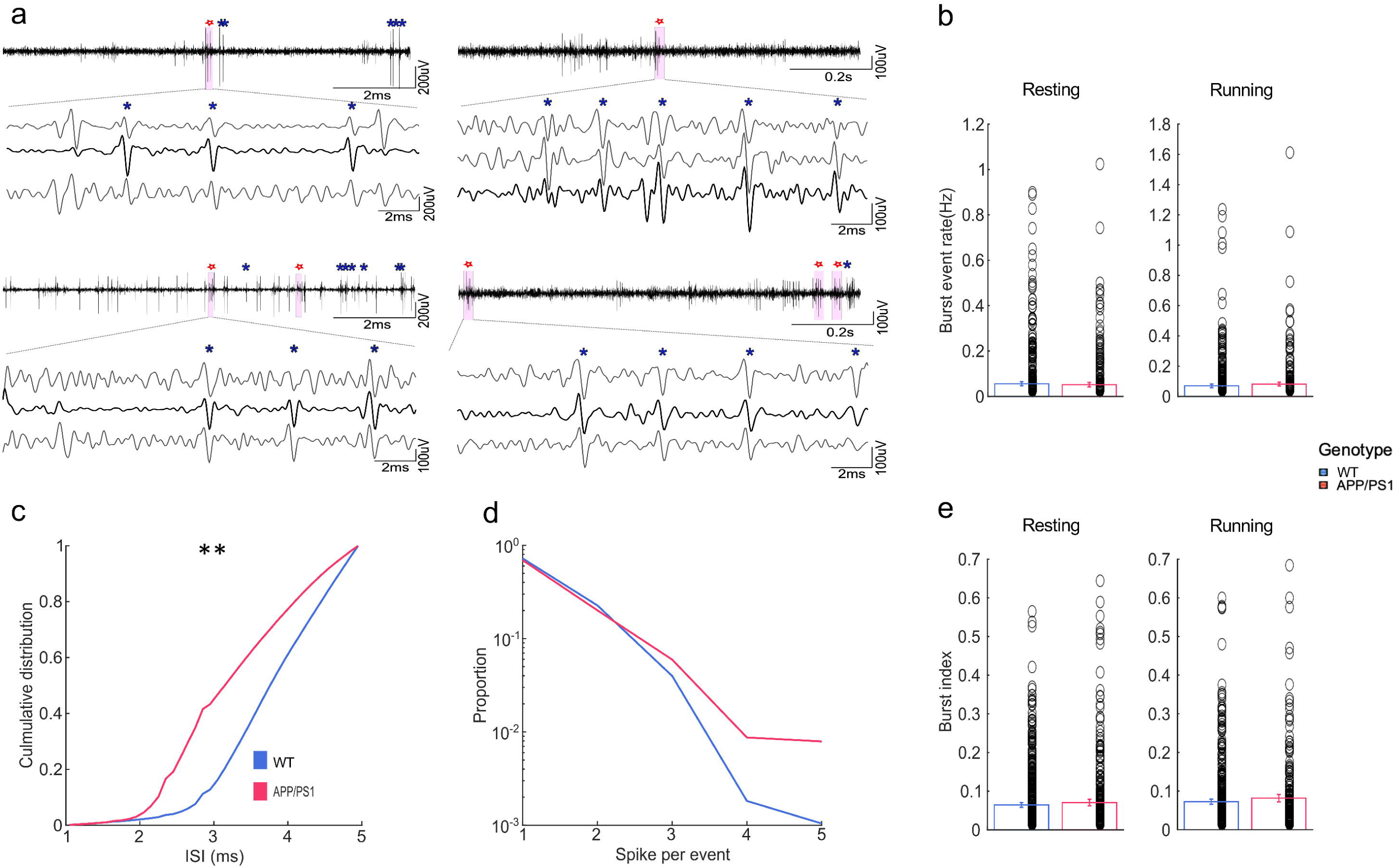
Bursting characteristics in awake behaving young APP/PS1 mice. (**A**) Representative burst events in high-pass filtered LFP recordings from WT (*left*) and APP/PS1(*right*) mice. Red stars indicate detected burst events, blue asterisks indicate single spikes. *Below*, zoomed-in burst events are shown, in three independent recording channels. (**B**) Burst event rate during resting and running. APP/PS1 mice exhibited a trend of higher burst event rate during running. (**C**) Cumulative distribution of inter-spike intervals (ISI) within burst events, showing significant shorter ISIs in APP/PS1 mice (*P* < 0.002, Kolmogorov–Smirnov test). (**D**) Distribution of spikes per burst, with APP/PS1 mice exhibiting more spikes per burst event. (**E**) Burst index during resting and running. APP/PS1 mice exhibited a trend of higher burst index during running. Asterisks indicate statistical significance (* *P* < 0.05, ** *P* < 0.01).

### Acceleration of hippocampal oscillations in awake behaving young APP/PS1 mice

Given our earlier findings of accelerated hippocampal oscillations in young APP/PS1 mice under general anaesthesia (Fig. 3), we next examined whether similar findings held in awake, drug-free conditions. To address this, we first analysed the power spectrum of hippocampal LFPs recorded during inter-trial periods in the linear track task, when animals briefly rested between task runs (Fig. 7a and b, Left). No significant difference was observed between APP/PS1 and WT mice in total power (Fig. 7b; 16887 ± 5407 µV Hz^-1^ for APP/PS1, n=7; 18376 ± 8931 µV Hz^-1^ for WT, n=9; *P* = 0.8015, f^2^(1,16) = 0.0657). However, APP/PS1 mice exhibited a significant increase of normalised power in beta band (15-25 Hz) (Fig. 7c; 0.0762 ± 0.0060 for APP/PS1, n=7; 0.0587 ± 0.0045 for WT, n=9; *P* = 0.0457, f^2^(1,16) = 4.8083), whereas the proportions of delta (0.1-4Hz, Fig7c; 0.2539 ± 0.0286 for APP/PS1, n=7; 0.2869 ± 0.0228 for WT, n=9; *P* = 0.2358, f^2^(1,16) = 1.5346), theta (4-12 Hz; Fig7d; 0.3632 ± 0.0317 for APP/PS1, n=7; 0.3416 ± 0.0211 for WT, n=9; *P* = 0.3315, f^2^(1,16) = 1.0120), and gamma (25-75 Hz; Fig7e; 0.0610 ± 0.0109 for APP/PS1, n=7; 0.0538 ± 0.0084 for WT, n=9; *P* = 0.7245, f^2^(1,16) = 0.1293) remained unchanged.

**Fig. 7.**
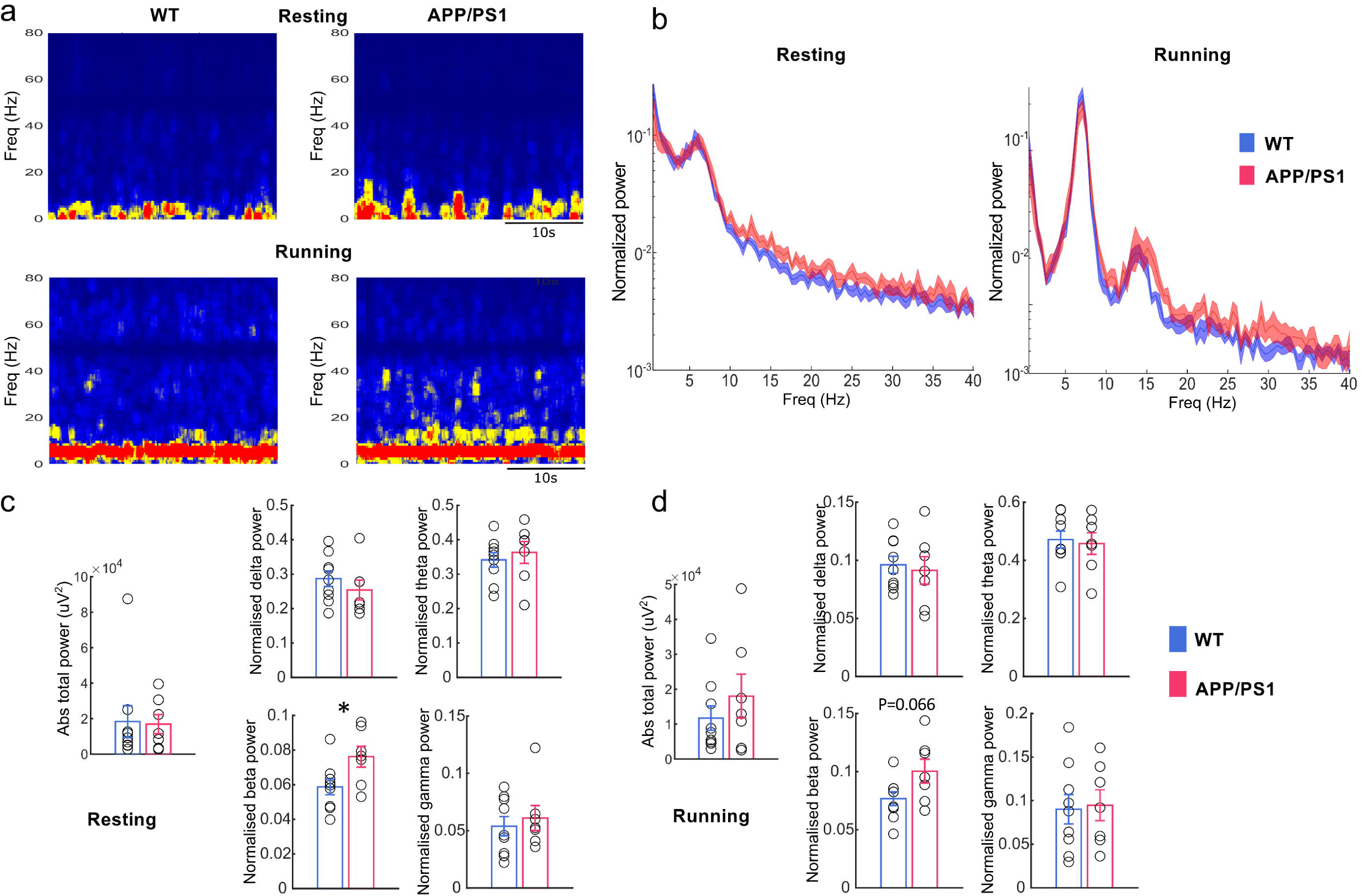
Characteristics of hippocampal oscillations in awake behaving young APP/PS1 mice. (**A**) Time-frequency representation of hippocampal LFP activity in WT and APP/PS1 mice during resting (left) and running conditions (right), respectively. The colour scale represents power intensity, with red indicating the highest power and blue indicating the lowest power. (**B**) Normalized power spectral density of hippocampal LFPs recorded during resting (left) and running (right) periods in WT and APP/PS1 mice. (**C**) Absolute total power and normalized power in different frequency bands (delta: 0.5–4 Hz, theta: 4–12 Hz, beta: 15–25 Hz, gamma: 40–80 Hz) during resting periods. APP/PS1 mice exhibited a significantly increased proportion of beta power compared to WT mice (*P* = 0.0457). (**D**) Same as (C) but during running periods. Asterisks indicate statistical significance (* *P* < 0.05, ** *P* < 0.01).

We further analysed power spectrum of hippocampal LFPs recorded when animals actively ran in the linear track task (Fig. 7a and b, Right). Again, no significant difference was observed between APP/PS1 and WT mice in total power during running (Fig. 7f; 18027 ± 6256 µV^2^ for APP/PS1, n=7; 11707 ± 3455 µV^2^ for WT, n=9; P =0.5784, f^2(^1,16) = 0.3237). However, APP/PS1 mice still exhibited an increase of normalised power in beta band (15-25 Hz), but not reaching the threshold for statistically significance (Fig. 7g; 0.1003 ± 0.0103 for APP/PS1, n=7; 0.0766 ± 0.0059 for WT, n=9; *P* =0.0657, f^2^(1,16) = 4.1525), whereas the proportions of delta (0.1-4Hz, Fig7h; 0.0913 ± 0.0119 for APP/PS1, n=7; 0.0962 ± 0.0072 for WT, n=9; *P* = 0.8015, f^2^(1,16) = 0.0657), theta (4-12 Hz; Fig7i; 0.4577 ± 0.0373 for APP/PS1, n=7; 0.4715 ± 0.0292 for WT, n=9; *P* = 0.8015, f^2^(1,16) = 0.0657), and gamma (25-75 Hz; Fig7j; 0.0947 ± 0.0178 for APP/PS1, n=7; 0.0901 ± 0.0170 for WT, n=9; *P* = 0.8015, f^2^(1,16) = 0.0657) remained unchanged. Together, LFPs recorded during both resting and running in awake animals reinforced the observation that young APP/PS1 mice exhibited a higher proportion of faster hippocampal oscillations.

## Discussion

This paper provides a quantitative analysis of the neurophysiological underpinnings of ‘early hippocampal hyperactivity’ *in vivo*, in a well-established amyloidopathy model of AD, the APP/PS1 mice. Our results indicate that at the single-cell level, 1) inhibitory synaptic input is reduced in pyramidal neurons but their intrinsic properties are largely unchanged; 2) pyramidal neurons show stronger bursts; 3) putative fast-spiking interneurons exhibit hypoactivity. At the neural network level, our data suggest: 1) a shift of hippocampal oscillation towards higher frequency ranges albeit no alteration in total oscillatory strength; 2) enhanced synchrony of synaptic excitation at higher oscillatory frequency and reduced overall oscillation-associated synaptic inhibition could explain accelerated hippocampal oscillations.

First, our findings directly reveal the cellular basis of ‘early hippocampal hyperactivity’ *in vivo* in early amyloidopathy. Hyperactivity may have simply implied that hippocampal neurons have an overall increased firing. Our results show this may not be the case. In contrast, excitatory pyramidal cells and inhibitory interneurons were differentially affected by the accumulation of soluble Aβ. Specifically, pyramidal cell intrinsic properties were largely unchanged, despite mildly narrowed action potentials. In awake, behaving animals, proportions of low, medium, and high firing rate pyramidal cells were largely unchanged, in agreement with a recent study using the same AD model in slightly older animals and miniature microscope Ca^2+^ imaging^27^. However, pyramidal cells did tend to fire stronger ‘bursts’, with a higher bursting frequency and more intra-burst spikes. These observations could at least partially explain previously observed ‘hyperactive’ phenotype in pyramidal neurons under two-photon Ca^2+^ imaging in young APP/PS1 mice^3^, in which the temporal resolution intrinsic to the kinetics of Ca^2+^ indicators may have hampered unequivocal discrimination between single spike and ‘burst’ of spikes. Previous findings from heavy Aβ plaque-bearing APP/PS1 mice also showed a profound increase of bursting in CA1 pyramidal cells *in vivo*^28^. Together with this finding, our data suggest the enhanced bursting in pyramidal cells could be a key cellular substrate of hippocampal hyperactivity in AD.

GABAergic interneurons have been suggested to be selectively vulnerable in AD^20,29–31^, but neurophysiological features of hippocampal GABAergic cells in AD *in vivo* are largely unclear, particularly at the early stage. Here, our results demonstrate a ‘hypoactivity’ phenotype in hippocampal fast-spiking interneurons in awake, behaving young APP/PS1 mice. Fast-spiking interneurons, which are typically parvalbumin (PV) positive, account for about 80% of the total interneuron population^32^. Given their peri-somatic axonal arborization into pyramidal cells, it is conceivable that a large proportion of inhibitory synaptic inputs recorded at the soma of pyramidal cells are of PV^+^ interneuron origin. Therefore, fast-spiking interneuron hypoactivity is consistent with the profound reduction of synaptic inhibition onto pyramidal cells observed in voltage-clamp recordings *in vivo*. Particularly, it explains the reduced frequency of sIPSCs seen in pyramidal cells, which suggests the involvement of a pre-synaptic mechanism, although the density of presynaptic interneuron populations (Supplementary fig. 2), inhibitory synapses (Supplementary fig. 3), and myelinated GABAergic axons (Supplementary fig. 4) were largely unaltered.

Second, our results shed light on the underpinning of ‘early hippocampal activity’ at the functional network level *in vivo* in early amyloidopathy. Compared to single-cell level alterations in neurophysiology, early markers at the functional network level may render higher translational value. Indeed, early hippocampal hyperactivity has been observed consistently in cognitively normal individuals who have a high-risk of developing AD, e.g. individuals of risk gene of familial AD^10,33^, cerebral Aβ deposition^34,35^, APOE4 carriers^10,13,14,36^, largely by fMRI scanning. Our data reveal that accelerated hippocampal oscillations may be able to bridge the gap between the enhanced BOLD signals and neural network activities that form the neural basis of key cognitive functions in the hippocampus, such as encoding and consolidating spatial memory. Our finding that the total strength of hippocampal oscillations is largely intact coincides with observations that young animals of sustained soluble Aβ do not show overt alterations in hippocampal related-cognitive functions^37^, and typical reduction of total neural activities only occurs at a later stage in people with clinical AD^8,38^.

Hippocampal slow waves such as delta oscillations serve as temporal framework for information transfer (e.g. memory) between the hippocampus and neocortex via coordinated activation of cell assemblies^39,40^. Given that impaired slow wave oscillations have been reported in people living with AD^41,42^ and in AD models^43,44^, the reduction of slow wave LFP components seen at the early stage of AD may suggest this key neural substrate for memory transfer is functionally deteriorating. Typically, in the hippocampus, rapid-eye-movement (REM) sleep is dominant by theta oscillations, whereas non-REM sleep is characterised by delta oscillations^45,46^. Although general anaesthesia is a distinct low-arousal state from sleep, reduced delta and increased theta in LFP power in our study could indicate a detrimental disruption of non-REM sleep in young APP/PS1 mice^27,44^, likely to reflect mild sleep fragmentation, a high-risk factor for clinical AD in humans^47^.

A recent study using the same animal model shows neocortical EEG power spectra shifts to higher frequencies at 12 months of age in both awake and sleep conditions^48^. The pattern and extent of neocortical EEG shifts across frequency ranges with heavy plaque burden in this study is very similar to our findings in the hippocampus where Aβ is still largely soluble. This contrast might suggest accelerated oscillations may not simply be a functional alteration restricted locally in the hippocampus but a progressing phenotype that may spread across brain regions with increasing Aβ burden. In addition, another study using awake APP/PS1 animals demonstrates a very similar increased relative power of beta oscillations and a general pattern of accelerated hippocampal oscillations across age^49^. Moreover, a recent human study targeting the hippocampus using magnetoencephalography (MEG) shows elevated faster oscillatory components in Aβ-bearing prodromal AD subjects^50^, corroborating our findings from the AD model. Taken together, accelerated hippocampal oscillations at early stage of AD may be considered as a translationally sensitive measurement which could indicate an early, yet sustained, and slowly spreading phenotype at the level of network function *in vivo*, a deviation from conventional view of oscillation slowing in AD. This might be particularly relevant due to the growing use of MEG in humans, which provides deep penetration and adequate temporal resolution for measuring hippocampal oscillations— something not achievable with EEG.

Finally, by exploiting simultaneous patch-clamp and LFP recording *in vivo*, we unravel the synaptic mechanism underlying accelerated hippocampal oscillations in early amyloidopathy. Oscillations typically reflect synchronous firings of neuronal assemblies^51^. For relatively slower oscillatory events such as delta, theta, and beta (e.g. from 0.1-25 Hz), their magnitude, measured by LFP power, depends largely on the strength of synchronized synaptic excitation (current generator), whereas synaptic inhibition typically provides a pace-making function in orchestrating synchronous firings of principal cells (rhythm generator)^52,53^. Within this framework, unchanged total power of oscillation-associated synaptic excitation may explain the largely unaltered total power of LFP. The reduced total power of oscillation-associated synaptic inhibition may well suggest an early deteriorating pace-making function for those oscillatory activities, which is yet to bring down the total level of orchestrated principal cell firing. In addition, this reduction of oscillation-associated inhibition is also in agreement with the overall reduction of basal synaptic transmission from *in vivo* voltage-clamp recordings as well as the ‘hypoactivity’ observed in fast-spiking GABAergic cell in awake, behaving animals. For specific oscillation components, the reduction of synaptic excitation power in the delta range and its enhancement in theta range coincide with the reduction of delta and the increase of theta oscillations in young APP/PS1 mice. Moreover, phase-locking of synaptic excitation is selectively enhanced to theta oscillations. Therefore, two lines of evidence indicate a stronger and more synchronous synaptic excitation of higher frequency is the main driving force for the accelerated LFP in early AD.

In summary, exploiting *in vivo* patch-clamp and high-throughput single-unit recordings, our results clarify the functional origins of early hippocampal hyperactivity in AD, shedding light on its underlying neurophysiological mechanisms at both cellular and network levels *in vivo*. This work resolves a longstanding uncertainty in the field. The observed acceleration of hippocampal oscillations may serve as a potential biomarker for early-stage AD and a therapeutic target for stabilizing neural networks, thereby preserving cognitive function before significant decline occurs.

## Data availability

Original data and scripts are available upon request from the corresponding author.

## Acknowledgements

We thank Prof. Tara Spires-Jones for sharing APP/PS1 mice, Dr Owen Dando for suggestions on statistical analysis, and Dr Sam Booker for advice on interneuron staining. We are grateful to Profs. Giles Hardingham, Tara Spires-Jones, and Matthew Nolan for helpful discussions and critical comments on the manuscript.

## Author contributions

SM performed *in vivo* patch-clamp and simultaneous LFP recordings. SK, ML performed high-throughput silicon probe recordings in awake animals with initial help from AK. SM, SK, and ML analysed data. XW helped with initial experiments and analysed data. MA, AT, LW, and CM contributed morphological and immunohistochemical experiments and analysis with inputs from LZ. JG helped with initial experiments. JG conceptualised and supervised all aspects of the project. JG wrote the paper with inputs from all authors.

