## supplementary material for "Reduced inhibition, bursting, and accelerated oscillations drive early hippocampal hyperactivity in Alzheimer’s disease"

### Supplementary materials

#### Extended materials and methods

##### *In vivo* patch-clamp and LFP recording

**Animals**: Male or female 2- to 3-month-old $APP^{swe}$/PS1ΔE9 mice and their littermate controls of C57BL/6J background were used. Mice were anesthetized by i.p. injection of 100 mg kg-1 ketamine (Rimadyl) and 10 mg kg-1 xylazine (Rompun), before mounting in a stereotaxic frame (David Kopf Instruments) and were supplied with 100% oxygen through a ventilation mask. Body temperature was continuously monitored by a rectal thermometer and maintained at 37 ± 0.5°C by placing the animal on a heating pad (Kent Scientific, UK).

**Surgery**: The skull of the animal was exposed and dried. A small craniotomy (~2 mm diameter) was made on the right or left hemisphere to target the dorsal hippocampus according to stereotaxic coordinates at AP ≈ 1.8 mm and ML ≈ 1.5 mm (AP, anterioposterior from bregma; L, lateral from midline). Subsequently, within the craniotomy window, the dura mater was carefully cut and removed. The exposed cortical surface was superfused with HEPES-buffered extracellular solution (135 mM NaCl, 3.5 mM KCl,1.8 mM $CaCl_2$, 1 mM $MgCl_2$, and 5 mM HEPES; pH = 7.28 with NaOH).

***in vivo* patch-clamp recording**: Patch pipettes were fabricated with a micropipette puller (P-1000; Sutter Instrument), using 1.0 mm / 0.5 mm (outer diameter / inner diameter) borosilicate glass tubing (Hilgenberg) and had tip resistance values of 4–6 MΩ. For voltage-clamp recordings, pipettes were filled with a $Cs^+$-based intracellular solution, containing 130 mM Cs-methanesulfonate, 2 mM KCl, 10 mM EGTA, 2 mM $MgCl_2$, 2 mM $Na_2ATP$,10 mM HEPES, 5 mM QX-314, and 3 mg ml-1 biocytin (pH adjusted to 7.28 with CsOH, 290–300 mOsm). For current-clamp recordings, pipettes were filled with an intracellular solution containing 130 mM K-methanesulfonate, 10 mM EGTA, 2 mM $MgCl_2$, 2 mM $Na_2ATP$, 10 mM HEPES, 2 mM KCl and 3 mg ml-1 biocytin (pH adjusted to 7.28 with KOH, 290–300 mOsm). A reference electrode (Ag–AgCl) was placed on the skull near the craniotomy window. The cortical surface was immersed with HEPES-buffered extracellular solution. The pyramidal cell layer of the dorsal hippocampus was targeted using age-corrected stereotaxic coordinates (AP: 1.8–2.0 mm; ML: 1.5–2.0 mm; and DV:1.1–1.4 mm; DV: dorsal-ventral from cortical surface). Patch pipettes were gently advanced from the craniotomy window perpendicular to the cortical surface with positive pressure (~300 mbar) applied to the pipette lumen to avoid tip plugging, until ~200 μm above the target area. Positive pressure was then reduced to ~15 mbar. Tight-seal cell-attached configuration was reached in the voltage-clamp mode, with pipette holding potential set at −70 mV to minimize holding current (typically 0 to –5 pA). After break-in, the whole-cell patch-clamp recording configuration was obtained, monitored by changes in current amplitudes in response to a 10-mV test pulse. Maximal care was taken to minimize the series resistance (Rs) during recording, which was 31 ± 1 MΩ in the present data set (range:12–50 MΩ). Rs was carefully monitored throughout the recording session using 20-ms,10-mV hyperpolarizing test pulses applied at ~1 min intervals. EPSCs and IPSCs were recorded in the voltage-clamp configuration with the same cell held at either −70 mV or +10 mV, respectively, with alternating order. Membrane potential values reported were not corrected for liquid junction potentials.

**Local field potential recording**: LFP recording pipettes were fabricated from 1.0 mm / 0.5 mm (outer diameter / inner diameter) borosilicate glass tubing (Hilgenberg) and had open-tip resistance values of 1–3 MΩ. LFP pipettes were coated with Dil before carefully inserted into the same craniotomy as the patch pipettes, at a 25° oblique angle, in the AP direction, targeting the CA1pyramidal cell layer of the dorsal hippocampus (AP: 1.8–2.0 mm, ML: 1.8–2.0 mm, DV:1.1–1.3 mm). Positive pressure (50–80 mbar) was applied to avoid pipette plugging. The location of LFP pipette was visualized by histology of the pipette track and by visualization of Dil. To unequivocally determine pipette location, only a single LFP pipette was inserted per animal.

**Biocytin labelling and cell visualization**: After recording, brains were fixed for >12 h in 4% paraformaldehyde, then stored in 0.1M phosphate buffer solution (PBS) at 4°C. To visualize labelled cells, the hemisphere containing the recorded cell was cut into 200-μm thick parasagittal sections using a vibratome (Leica VT 1000S, Germany), washed in 0.1 M PBS containing 0.3% Triton-X and 0.05% sodium azide (PBS-T-A), and then incubated with streptavidin Alexa

Fluor® 488 (1:1000 in PBS-T-A; ThermoFisher, UK) at 4°C overnight. Sections were then washed in 0.1 M PBS, rinsed in dH2O, and mounted using VectaShield (Vector Labs, UK). Sections were imaged using a confocal microscope (SP8, Leica, Germany).

**Data acquisition**: *in vivo* patch-clamp and LFP recordings were made using an EPC 10 Quadro amplifier (HEKA, Germany) and InstruTech LIH 8+8 data acquisition system (HEKA, Germany). Both signals were low pass-filtered at 10 kHz (Bessel), sampled at 20 kHz, and stored using Patchmaster v2x91 software running on a PC under Windows 10.

**Synaptic currents**: Synaptic charge was calculated as the area under the curve (baseline normalized to 0, area under the curve calculated using trapz in MATLAB) from a 30 second recording sweep at either -70 mV or +10 mV for excitation or inhibition respectively. E/I balance was calculated as excitation charge/inhibition charge. For the quantification of spontaneous excitatory postsynaptic currents (sEPSC) and spontaneous inhibitory postsynaptic currents (sIPSC), first, the dv/dt of the trace was calculated, and peaks were detected when exceeding a set threshold of (minimum of 1.8 for EPSC and 1.2 for IPSC, with threshold varying based on signal noise and activity level) standard deviations above the mean. A minimum amplitude threshold was set for each of these events which varied based on signal noise (minimum of 7 pA for excitation and minimum 15 pA for inhibition). To select only individual PSC events, and not cumulative events, the start of the PSC was then selected if it was within a set number of standard deviations (varied based on levels of noise and activity, minimum of 1.5 for EPSC, 0.3 for IPSC) from baseline. The end current detected also had to be within a set range of the start value (minimum of 10 pA and 70 pA for EPSC and IPSC respective but varied depending on signal) to eliminate multiple cumulative events and detected events had to be a minimum distance apart (200 and 100 samples for EPSC and IPSC respectively but depending on signal noise and activity). From this, detected events were extracted and summarized. Average waveforms were extracted and used to calculate the mean amplitude, the rate of rise, and decay tau (fitted on 20-80% of the curve). Frequency was taken as the number of individual events, normalized per second in a 30 second sweep. The interevent interval between events was also calculated.

**Passive and active properties**: Resting membrane potential (Vm) was taken as the membrane potential immediately after break-in. Passive and active properties including excitability measures were calculated from the current step injections. From the -100pA hyperpolarizing sweep, input resistance was calculated, based on the difference from Vm just prior to current injection compared to the average at steady state at the end of the current injection. A dv/dt and amplitude threshold was set to detect action potentials. From detected action potentials, the maximum dv/dt rate, action potential amplitude, peak value, threshold, width at half height and width at threshold were calculated. Input- frequency plots were made based on the number of quantified action potentials per current injection sweep.

**Power spectrum density calculation and power analysis**: Data analysis was performed using custom-made scripts in Matlab (R2024b, Mathworks). Mains power interference (50,138,150,250 Hz) was first removed from LFP and synaptic current (EPSCs & IPSCs) signals by a second order band-stop filter. Signals were down-sampled to 2 kHz, using re-sample function which applied a FIP anti-alising low-pass filter, and then filtered using a 4th-order Butterworth band-pass filter between 0.1 Hz and 300 Hz. LFPs with large baseline drift or clear noise were removed. Slow drifts in synaptic currents were baseline-corrected by subtracting their respective means. Power spectrum was obtained by pwelch function (window: 2s, overlap: 50%). Mains power interference (50&138&150&250Hz) still appearing in the power density data was amended by replacement with the mean of neighbouring density values. Absolute power was calculated as the sum of power spectral density within each frequency band. Normalized power was calculated from dividing absolute power by total power (0.1~300Hz).

**Derivative-based detection of synaptic events and phase analysis**: To quantitatively examine the temporal structure and relationship between synaptic currents and hippocampal oscillations *in vivo*, we used a first derivative-based detection method. Synaptic currents were smoothed using a moving average filter with a 2.5 ms time constant. Next, we calculated the first derivatives of the smoothed traces. EPSC onsets were detected as derivative minima, and IPSC onsets as derivative maxima. To account for variability in synaptic current kinetics, we analyzed subsets of data corresponding to the largest derivative peaks (top 1–25% percentiles). To calculate phase angles, wide-band LFP signals were filtered between 0.1–4 Hz for delta and 4–12

Hz for theta oscillations. The Hilbert transform was then applied to the band-pass-filtered LFP signals. Next, each EPSC or IPSC onset time point was assigned a Hilbert phase value. The mean phase angle was then obtained for each cell and averaged across cells. As an indicator of the locking quality of EPSC or IPSC phases, we used vector strength. Vector strength was calculated for different subsets of data representing the largest derivative peaks (top 1–25% percentiles), and then averaged across these subsets. Rayleigh test was used to assess non-uniformity of phase distributions.

#### High-throughput silicon probe recordings in awake, behaving animals

**Surgical Preparation**: Male or female 3- to 5-month-old $APP^{swe/}PS1\Delta E9$ mice and their littermate controls of C57BL/6J background were used. Mice were anesthetized with isoflurane (3% induction, 1% maintenance, 1 L/min in pure oxygen) and secured in a stereotaxic frame. Two steel screws (1 mm in diameter) were placed in small boreholes drilled above the cerebellum and olfactory bulb. A custom-made head-ring for subsequent head-restraint training and recording was attached to the skull, with the screws firmly fixed using bone cement (Refobacin, Biomet). Before surgery, mice received a subcutaneous injection of carprofen (120–130 µL, 20 mg/kg) and an subcutaneous supplement of 0.9% NaCl (200–300 µL). Mice were given at least seven days to recover before further procedures. After recovery, mice were handled and habituated to head-restraint on an air-cushioned styrofoam ball in a virtual reality (VR) system (JetBall, Phenosys). Following habituation, mice were trained to run in a VR linear treadmill task for liquid rewards. Once the mice demonstrated consistent running behavior after 3–4 days, they were transiently anesthetized, two small craniotomies were drilled above the hippocampus in each hemisphere (AP: ≈−1.9 mm; ML: ≈±1.5 mm). The dura was carefully removed, and the craniotomies were sealed with silicone (Kwik-Cast, World Precision Instruments). Mice were allowed to recover for at least eight hours before electrophysiological recordings began.

**VR Linear Treadmill Task**: For the VR linear treadmill task, head-fixed mice were positioned on an air-cushioned styrofoam ball within a 270° TFT surround monitor system (JetBall, PhenoSys). Mice were trained to run through a custom-designed virtual linear corridor, covering a 150 cm running path per trial. Upon reaching the end of the corridor, they received a 50µL liquid reward (5% sucrose water) via a spout. After a 10-second inter-trial interval, mice were teleported back to the start of the corridor to begin the next trial. Each training session lasted 15–30 minutes daily for at least three days. Trial events were recorded as transistor-transistor logic (TTL) pulses by the acquisition device, while locomotion data were collected using an XY-motion sensor at 50 Hz.

**Silicon probe recordings**: Mice were secured in the stereotaxic frame within the VR system and allowed to perform the linear treadmill task. Silicon probe (A4X32-Poly2–5mm-23s-200-177, NeuroNexus), containing 128 recording channels across four parallel shanks, was slowly inserted into the hippocampus through the previously prepared craniotomies. Neural signals were recorded using an Intan 128-channel head-stage (C3316, Intan Technologies) connected to an Intan RHD recording controller (C3004, Intan Technologies). Signals were sampled at 20 kHz and digitized as 16-bit signed integers.

**Spike sorting and classification of cell types**: Spike sorting was performed using Kilosort 3, followed by manual curation in Phy 2. Spike clusters were assigned to the channel with the largest voltage trough-to-peak amplitude, determined from the average spike waveform. Only units displaying a clear refractory period (± 1.5 ms) were included in the further analysis. To classify single-unit spike clusters into putative excitatory and inhibitory neurons, single-units were distinguished based on spike waveform shape and the first moment of the autocorrelogram. First, spike width was measured by calculating the time between trough and peak from the average spike waveform of each cluster. Second, the first moment of the autocorrelogram was calculated as the center of mass along the time axis of an autocorrelogram computed with lags from 0 to 50 ms. Single units were excluded if their autocorrelograms had a peak spike count of less than 10. Putative pyramidal cells were defined as units with a spike width greater than 0.5 ms and a first moment of the autocorrelogram less than 25 ms. Putative interneurons were defined as units with a spike width less than 0.5 ms and a first moment of the autocorrelogram greater than 20 ms.

**Identification of burst firing**: Complex bursts were classified as a series of three or more spikes with inter-spike intervals of less than 5 ms. Burst event rate was calculated as the number of burst events divided by the total recording time. The burst index was defined as the ratio of bursting spikes to all spikes.

**LFP analysis**: LFP signals were obtained by downsampling raw traces to 2 kHz and bandpass filtering between 0.1 and 300 Hz. LFP signals from each site were bandpass filtered (150–250 Hz), and power was calculated using the Hilbert transform. The channel with the largest mean power of filtered LFP was determined for each shank in each session and designated as the pyramidal cell layer. The power spectral density of neural activity in the pyramidal layer was computed using the Matlab function pwelch (window: 2s; overlap: 50%). Power was calculated as the area under the curve (AUC) of the power spectral density within each frequency band: delta (0.1 ~4Hz), theta (4–12 Hz), beta (15–25 Hz), and gamma (25–75 Hz). Normalized power was calculated as each frequency power divided by the total power. LFP was further analyzed to identify theta for 'running' and non-theta 'resting' periods. Signals were bandpass filtered for delta (2–4 Hz) and theta (4–10 Hz) frequency bands. Root mean square was calculated in each filtered LFP and the theta/delta power ratio was measured in 1600 ms segments with 800 ms overlapping steps. Theta epochs were defined as periods where the theta/delta ratio exceeded 2 for at least 2.4 seconds. Non-theta epochs were selected when the theta/delta ratio dropped below 2 for at least 4 seconds.

#### Immunofluorescence staining of interneurons, synapses, myelination, and amyloid plaques

Following transcardial perfusion, brains were fixed in 10% formalin for 2 hours and subsequently washed 3 times in 1X phosphate buffered saline (PBS). Brains were then dehydrated in 20% sucrose solution for 48 hours, snap-frozen in isopentane and stored in at -80 ºC. Brains were sectioned into 40 µm thick slices with a cryostat machine (CM1950, Leica Microsystems), which were kept at -20 °C in anti-freezing media (500ml 1X PBS, 85.6g sucrose, 1.42g magnesium chloride hexahydrate, filled to 1 L with glycerol) until immunostaining. Sagittal mouse hippocampal brain sections from either left or right hemisphere spaced 480 µm apart were used for each staining condition. For each brain, 3 technical replicates were used. The following primary antibodies were used: parvalbumin (rabbit, PV27a, 1:1000, Swant), somatostatin (rabbit, T-4103, 1:1000, BMA Biomedicals), VGAT (guinea pig, 131004, 1:250,

Synaptic Systems), gephyrin (mouse, 147021, 1:100, Synaptic Systems), MBP (rat, MCA4095, 1:300, BioRad), β-amyloid 6E10 (mouse, 803004, 1:1000, BioLegend). The secondary antibodies were as follows: Alexa Fluor 594 goat anti-mouse (A11005, 1:1000, Thermo Fisher Scientific), Alexa Fluor 488 goat anti-guinea pig (A11073, 1:1000, Thermo Fisher Scientific), Alexa Fluor 594 goat anti-rat (A11007, 1:1000, Thermo Fisher Scientific), Alexa Fluor 488 goat anti-mouse (A21121, 1:1000, Thermo Fisher Scientific), Alexa Fluor 488 goat anti-rabbit (A11008, 1:500, Thermo Fisher Scientific). The immunostaining protocols used in this study are summarised below.

**Inhibitory interneuron staining**

Sections were washed 3 times in 1XPBS and blocked for 1 hour (10 % normal goat serum (NGS), 0.3 % Triton X-100, 0.05 % sodium azide in PBS). Sections were incubated with primary antibodies in a blocking solution (5 % normal goat serum, 0.3 % Triton X-100, 0.05 % sodium azide in PBS) for two days at 4 °C. Next, sections were washed 4 times in 1XPBS and incubated with the secondary antibodies in blocking solution (3 % normal goat serum, 0.3 % Triton X-100, 0.05 % sodium azide in PBS) overnight at 4 °C. The following day, sections were washed 3 times in 1XPBS, then 2 times in 0.1 M phosphate buffer (PB). Nuclei were stained with 4′,6-diamidino-2-phenylindole (DAPI) in PB for 1 hour at room temperature.

**Myelin basic protein (MBP) and PV immunostaining**

Sections were washed 2 times with 1XPBS, then blocked in blocking solution (10 % NGS in PBS-Triton X-100 0.2 %) for 2 hours. Primary antibodies were diluted in blocking solution and sections were incubated overnight at 4 °C. The next day, sections were washed 3 times in PBS-Triton X-100 0.2 % and incubated with the secondary antibodies in blocking solution for 2 hours at room temperature. Lastly, sections were washed once in PBS-Triton X-100 0.2 %, counterstained with DAPI, and washed again 3 times with 1XPBS.

**Synaptic staining**

Sections were washed in 1XPBS, followed by antigen retrieval with a citrate-based antigen unmasking solution (H-3300-250, 1:100, Vector Laboratories) at 95 °C for 20 minutes. Sections were first washed once in 1XPBS, then once in PBS-Triton X-100 0.3 % (PBT), and were incubated in Image-iT® FX Signal Enhancer (I36933, Thermo Fisher Scientific) for 30 minutes. Sections were washed once in 1XPBS and blocked with blocking solution (10 % heat-inactivated horse serum, 0.3 % Triton X-100, 1X PBS) for 2 hours. Primary antibodies were diluted in blocking solution and sections were incubated for two days at 4 °C. Next, sections were washed 4 times in PBT (1h/wash) and subsequently incubated with secondary antibodies in blocking solution overnight at 4 °C. The following day, sections were washed 3 times in PBT, counterstained with DAPI, and washed again once in PBT.

**Amyloid plaque staining**

Sections were washed twice in PBS, then blocked for 2 hours with blocking solution (10 % normal goat serum, 0.2 % Triton X-100, 1X PBS). Primary antibodies were diluted in blocking solution and brain sections were incubated overnight at 4 °C. The following day, sections were washed 3 times in PBST (0.2 % Triton X-100, 1X PBS), then incubated with the secondary antibody diluted in blocking solution for 2 hours. Finally, sections were washed once in PBST, then counterstained with DAPI, and washed once with PBS.

All sections were mounted onto microscopic slides using ProLong Gold Antifade Mountant (P36930, Thermo Fisher Scientific).

#### Image acquisition and analysis

All images were acquired with a confocal microscope (Carl Zeiss LSM 800 with AiryScan 2, Germany), using 10X, 20X, or 63X objectives. Image processing and analysis was done with the ZEN Microscopy Software, ImageJ and Fiji software, and MATLAB.
Quantitative data are expressed as mean ± standard error of the mean (SEM). Normality of data was assessed by Shapiro-Wilk test, and parametric or non-parametric analysis was selected accordingly. Difference between groups was analysed using two-way ANOVA to account for sex and age differences, or unpaired t-test. The significance level for analysis was set to $p<0.05$. All statistical analysis was performed using the software GraphPad Prism (GraphPad Software, USA).

**Quantification of parvalbumin and somatostatin-expressing interneurons**

The hippocampal CA1 was traced using the 20X/0.45 objective and Z-stacks spaced 0.85 µm apart. Positively stained interneurons in the CA1 were quantified. Image analysis was performed using ZEN Microscopy Software. The total number of PV+ and SST+ neurons was determined manually, with neurons only being included in the quantification if they had a clearly stained soma and dendrites. Final cell density was determined by dividing the number of quantified positively stained cells by the tissue volume (area of the hippocampus multiplied by section thickness).

**Measurement of MBP and PV colocalization**

Images of the CA1 were captured using a 20X objective and Z-stacks (0.85 µm interval between images) and reconstructed with maximum intensity projection. Tile scanning was used to acquire images covering the CA1 area of the hippocampus. Analysis was performed using ImageJ and Fiji as previously described. Automated thresholding using Otsu's method was performed on the output images. Immunoreactivity for MBP and PV were determined as the number of signal-positive pixels in the thresholded images. Overlap between the positive MBP and PV signals was determined using the image calculator function. For MBP colocalisation to PV-expressing interneurons, the number of MBP+ and PV+ pixels were quantified and expressed as a percentage of total PV signal-positive pixels.

**Quantification of synapses**

Images covering the pyramidal layer were captured using a 63X objective and Z-stacks (acquired at 0.2 µm intervals). Representative images of the CA1 region were captured with a 63X objective. Acquired images were divided into 10 µm x 10 µm regions of interest, followed by cropping and segmentation using automated thresholding in ImageJ. Pre- and post-synaptic objects were quantified with MATLAB using a custom script. Obtained values were expressed as synapses/mm3 per CA1 area (pyramidal and dendritic) and averaged for each brain.

**Quantification of amyloid plaques**

Images covering the whole brain section were captured using a 10X objective and Z-stacks (acquired at 1.50 µm intervals). Positively stained amyloid plaques in the hippocampus and neocortex were quantified. Image analysis was performed using ZEN Microscopy Software. The total number of amyloid plaques was determined manually. Final plaque density was determined by dividing the number of quantified amyloid plaques by the tissue volume (area of the hippocampus multiplied by section thickness).

### Supplementary figures

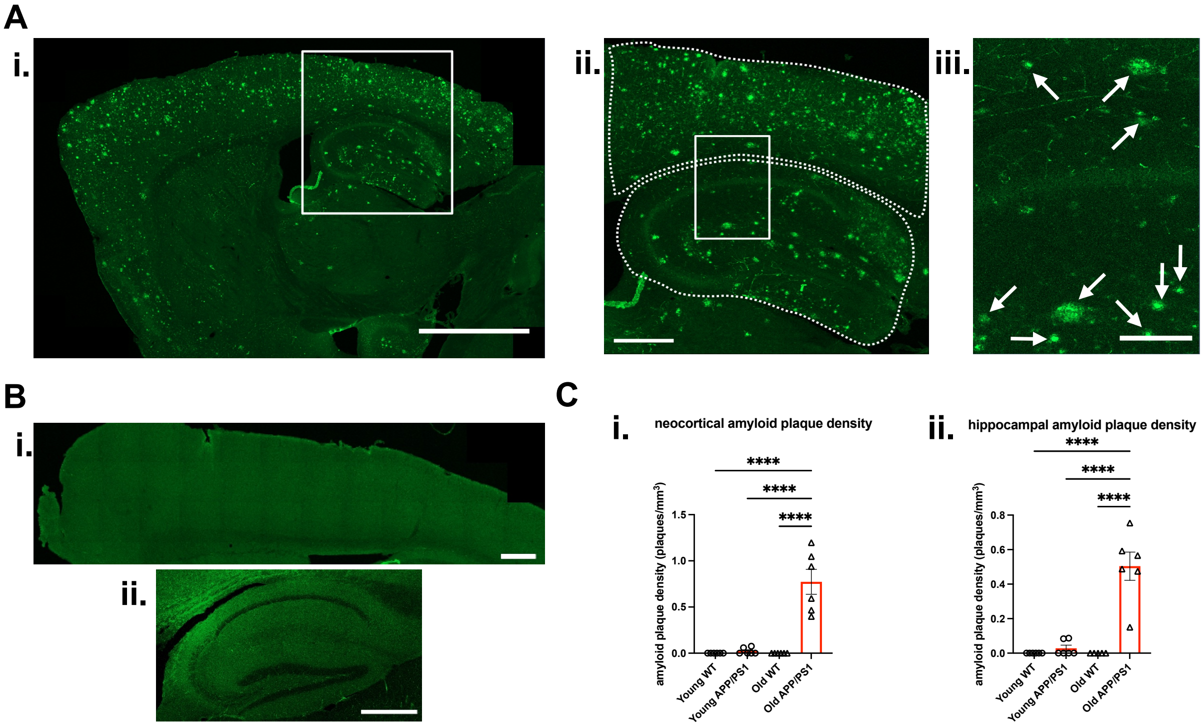


**Supplementary fig.1 Amyloid plaque accumulation in the hippocampus and neocortex in young (3-5.5 months) and old (11-13 months) APP/ PS1 mice.**
**A.** Representative images of amyloid plaque accumulation in old APP/PS1 mice. Scale bar: 2000µm. **ii.** Inset image of framed area in (i.) showing an example of the surface areas used for quantification surrounded by a dotted line (top: neocortex, bottom: hippocampus). Scale bar: 500µm. **iii.** Inset image of framed area in (ii.) with arrows indicating amyloid plaques. Scale bar: 200µm. **B.** Representative image of absence of amyloid plaque accumulation in young APP/PS1 mice in the **i.** neocortex and **ii.** hippocampus. Scale bar: 500µm. C. Data showing amyloid plaque density in the **i.** neocortex and **ii.** hippocampus. Density was expressed as number of plaques per $mm^3$. Statistical analysis was done with a two-way ANOVA. Error bars represent mean ± SEM, n number represents number of mice used, **** $p<0.0001$.

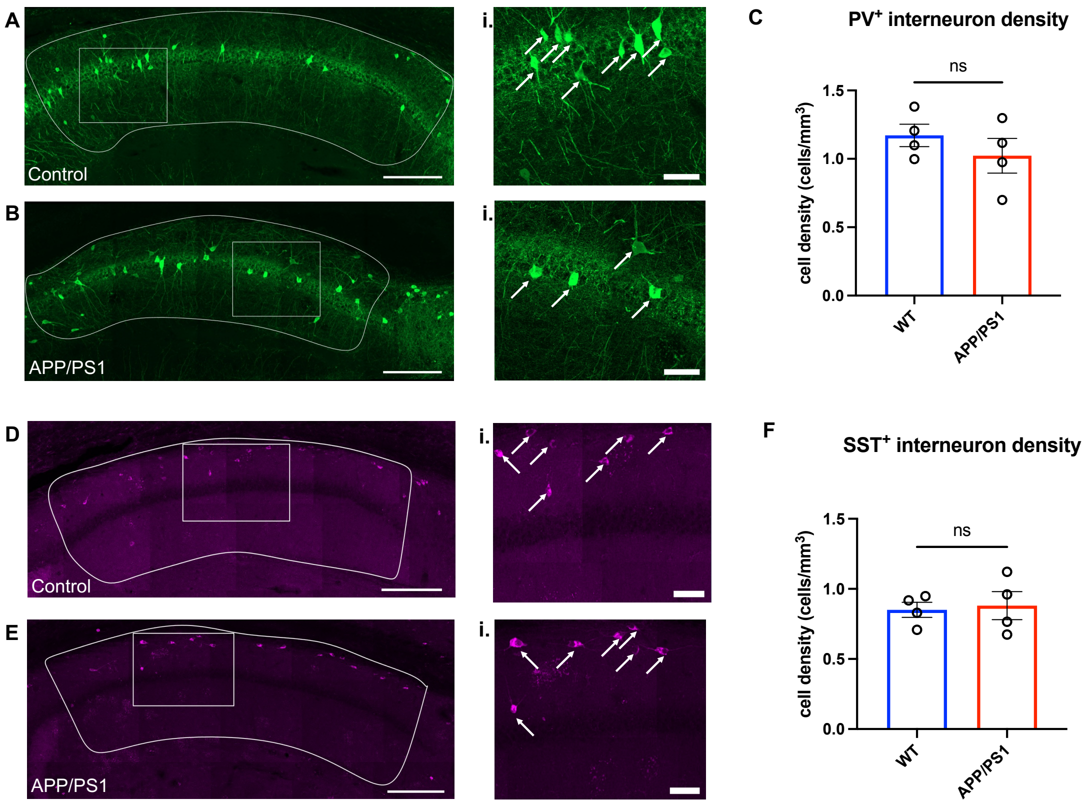


**Supplementary fig.2 Parvalbumin (PV)$^+$ and somatostatin (SST)$^+$ interneuron density in the hippocampal CA1 region of young mice.**

**A-B.** Representative images showing immunofluorescence staining of PV$^+$ interneurons in **A.** control and **B.** APP/PS1 hippocampi of young mice. Scale bar: 200 µm. Manually surrounded area represents CA1 hippocampi area used for PV$^+$ interneuron density quantification. **i.** Enlarged representative image of the CA1 area of the hippocampus of boxed areas in A and B. Arrows indicate positively stained neurons. Scale bar: 50 µm. **C.** Data showing cell density for PV$^+$ interneurons in the hippocampal CA1 region of young mice. Comparisons were done with an unpaired t test. Error bars represent mean ± SEM, n number represents number of mice used, ns: not significant. **D-E.** Representative images showing immunofluorescence staining of SST$^+$ interneurons in **D.** control and **E.** APP/PS1 hippocampi of young mice. Scale bar: 200 µm. Manually surrounded area represents CA1 hippocampi area used for PV interneuron density quantification. **i.** Enlarged representative image of the CA1 area of the hippocampus of boxed areas in A and B. Arrows indicate positively stained neurons. Scale bar: 50 µm. **F.** Data showing cell density for SST$^+$ interneurons in the hippocampal CA1 region of young mice. Comparisons were done with an unpaired t test. Error bars represent mean ± SEM, n number represents number of mice used, ns: not significant.

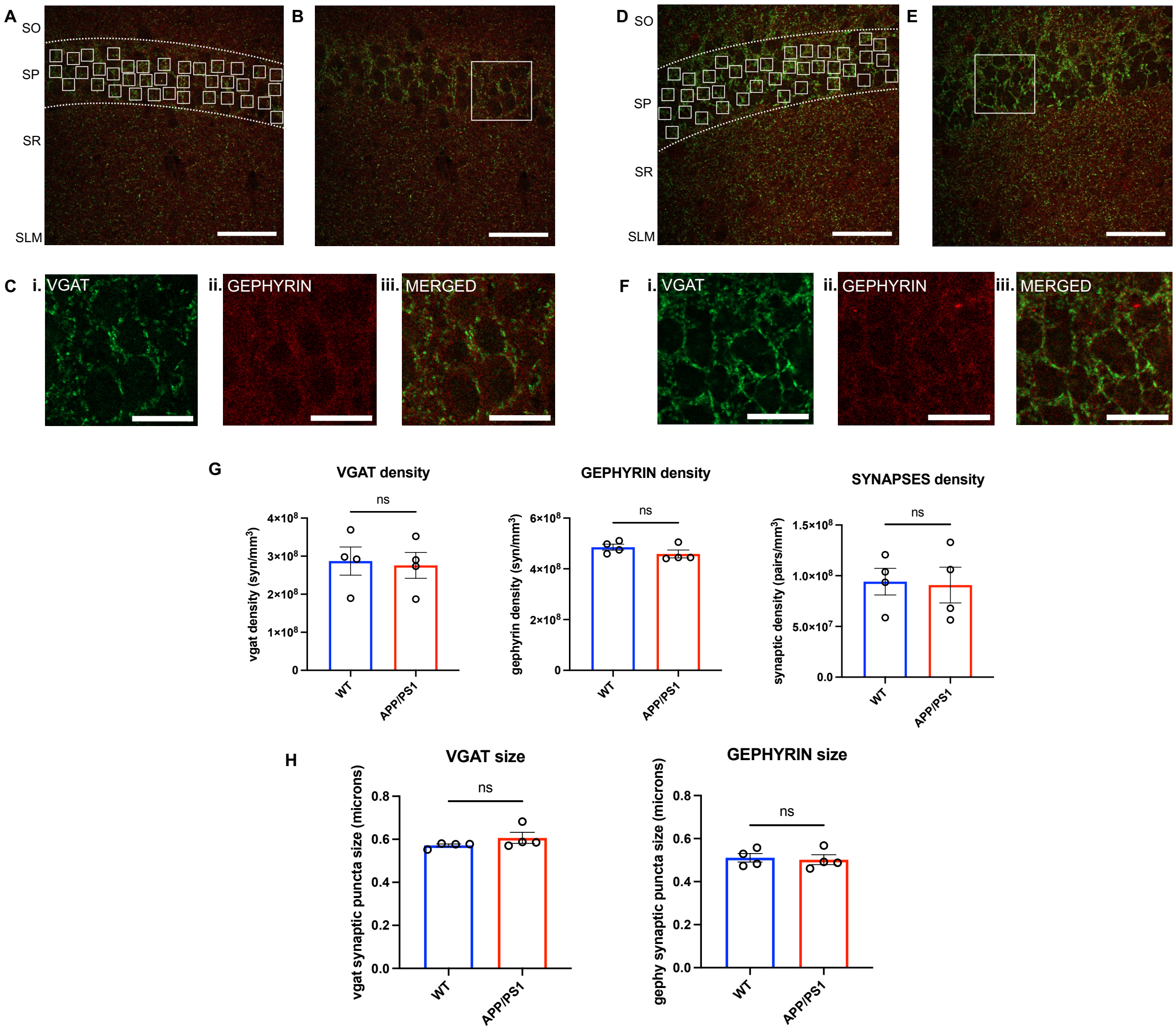


**Supplementary fig.3 The quantification of perisomatic inhibitory synapses in the hippocampal CA1 region of young mice.**

**A.** Representative immunostaining of vgat and gephyrin expression in the hippocampal CA1 of young WT mouse, with white regions of interest (ROIs) (10x10μm) indicating regions used for perisomatic inhibitory synapse quantification. Scale bar: 50 μm. **B-C.** Enlarged view of boxed area in panel B showing **i.** vgat stain (perisomatic basket shape pattern observed), **ii.** gephyrin stain and **iii.** synaptic pairs (vgat and gephyrin merged). Scale bar: 20um. **D.** Representative immunostaining of vgat and gephyrin expression in the hippocampal CA1 of young APP/PS1 mouse, with white regions of interest (ROIs) (10x10μm) indicating regions used for perisomatic inhibitory synapse quantification. Scale bar: 50 μm. **E-F.** Enlarged view of boxed area in panel E showing **i.** vgat stain (perisomatic basket shape pattern observed), **ii.** gephyrin stain and **iii.** synaptic pairs (vgat and gephyrin merged). Scale bar: 20um. **G.** Data showing quantification of vgat, gephyrin (synapses/$mm^3$) and synaptic pairs density (pairs/$mm^3$) respectively. The brain region quantified was the pyramidal layer of the hippocampal layer CA1, n number represents number of mice used. Statistical analysis was done using unpaired t test. Error bars represent mean ± SEM, ns: non-significant. **H.** Data showing synaptic puncta size in microns for vgat and gephyrin synapses, respectively. The brain region quantified was the pyramidal layer of the hippocampal layer CA1, n number represents number of mice used. Statistical analysis was done using unpaired t test. Error bars represent mean ± SEM, ns: non-significant.

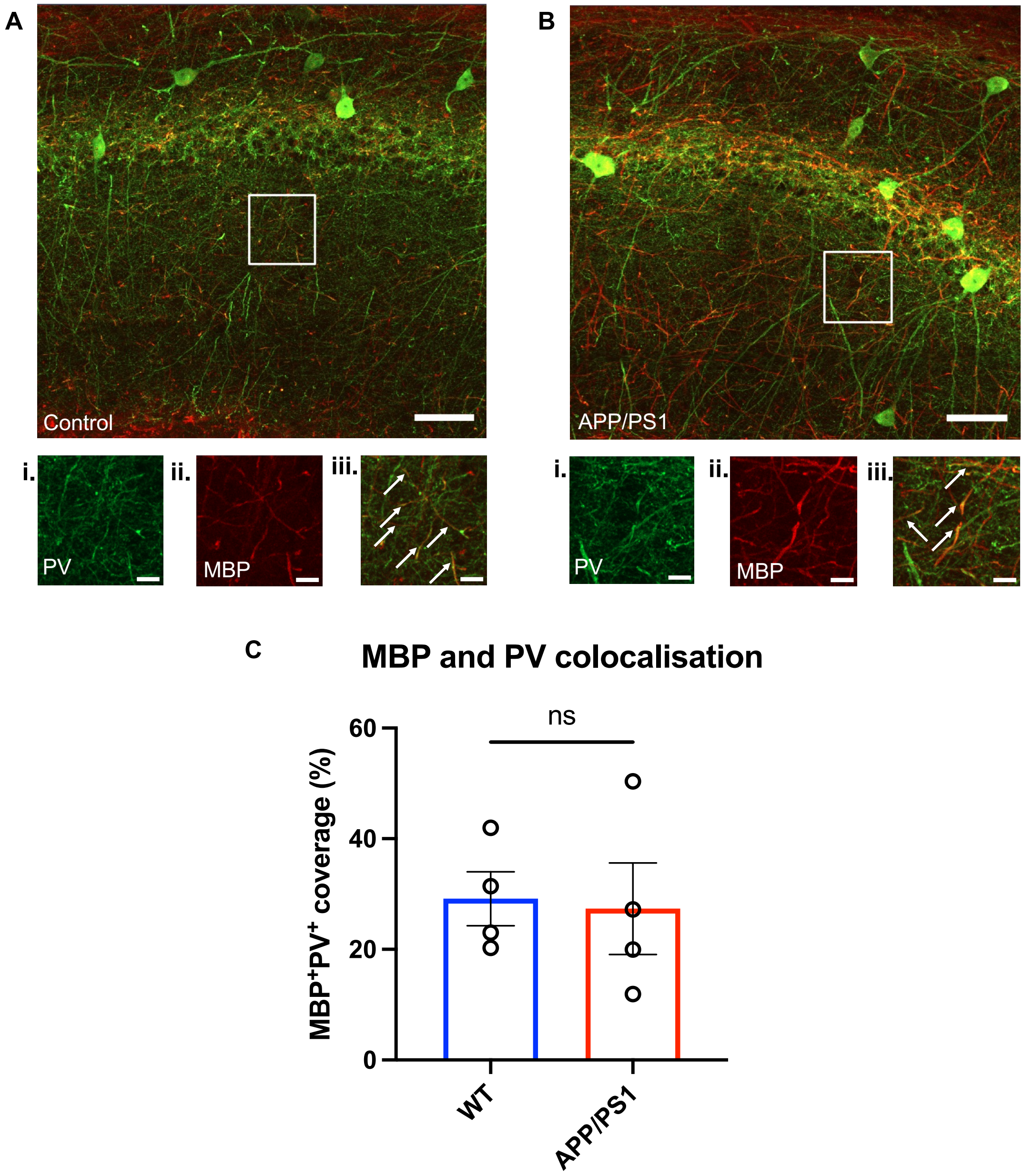


**Supplementary fig.4 Myelination of the hippocampal CA1 parvalbumin-positive interneurons in young mice. A-B.** Representative images of WT (A) and APP/PS1 (B) CA1 region showing immunoreactivity for parvalbumin (PV) and myelin basic protein (MBP) in young mice. Scale bar: 50 µm. Inset images of the frames areas in A and B show single channels of i. PV and ii. MBP and iii. merged channels. Arrows in iii. indicate overlap of PV and MBP immunoreactivity. Scale bar represents 10 µm **C.** Data showing relative colocalization of MBP and PV in the CA1 of young WT and APP/PS1 mice. $PV^+$ and $MBP^+$ area was expressed as a percentage of the total $PV^+$ area. Statistical analysis was done with an unpaired t test. Error bars represent mean ± SEM, n number represents number of mice used, ns: not significant.
